# Chromatin binding by HORMAD proteins regulates meiotic recombination initiation

**DOI:** 10.1101/2023.03.04.531117

**Authors:** Carolyn R. Milano, Sarah N. Ur, Yajie Gu, Eelco C. Tromer, Jessie Zhang, Andreas Hochwagen, Kevin D. Corbett

## Abstract

The meiotic chromosome axis coordinates chromosome organization and interhomolog recombination in meiotic prophase and is essential for fertility. In *S. cerevisiae*, the HORMAD protein Hop1 mediates enrichment of axis proteins at nucleosome-rich genomic islands through a central chromatin-binding region (CBR). Here, we use cryoelectron microscopy to show that the Hop1 CBR directly recognizes bent nucleosomal DNA through a composite interface in its PHD and winged helix-turn-helix domains. Targeted disruption of the Hop1 CBR-nucleosome interface causes loss of axis proteins from nucleosome-rich islands, reduces meiotic DNA double-strand breaks (DSBs), and leads to defects in chromosome synapsis. Synthetic effects with the disassemblase Pch2 suggest that nucleosome binding delays a conformational switch in Hop1 from a DSB-promoting, Pch2-inaccessible state to a DSB-inactive, Pch2-accessible state to regulate the extent of meiotic DSB formation. Phylogenetic analyses of meiotic HORMADs reveal an ancient origin of this domain, suggesting that these mechanisms are broadly conserved.

## Introduction

Across eukaryotes, sexual reproduction requires meiosis, a two-stage cell division program that produces haploid gametes from a diploid progenitor cell. The reduction of ploidy requires recombination of homologous chromosomes to form interhomolog crossovers (COs), which facilitate chromosome alignment on the meiosis I spindle. COs result from the repair of programmed DNA double-strand breaks (DSBs) that are introduced throughout the genome at the onset of meiosis. Failure to form sufficient DSBs or interhomolog COs can lead to errors in chromosome segregation and aneuploidy ^1,2^.

Meiotic recombination occurs in the context of tightly choreographed changes in chromosome morphology that regulate the number and position of DSBs and CO repair events. These morphology changes are mediated in large part by proteins of the meiotic chromosome axis, or axial element. Axis proteins bind replicated chromosomes to organize each chromosome into a linear array of chromatin loops and help activate, distribute, and resolve meiotic recombination events ^3–5^. The axis comprises three main components: (1) cohesin complexes with one or more meiosis-specific subunits ^6^; (2) filamentous axis core proteins 7; and (3) meiotic HORMA domain (HORMAD) proteins ^8,9^. Cohesins are ring-like complexes that bind DNA and processively extrude chromatin loops to organize chromosomes throughout the cell cycle 10. Cohesin complexes bind the filamentous axis core proteins, mediating the assembly of extended coiled-coil filaments that anchor DNA loops ^7^. Once assembled, the filamentous core proteins recruit HORMADs which serve as master regulators of meiotic recombination and promote the stable alignment of recombining chromosomes within the synaptonemal complex ^11,12^.

The hierarchical assembly of axis proteins, as well as the function of HORMAD proteins in the activation and later repair of meiotic recombination, is well conserved across eukaryotes, but is perhaps best understood in the budding yeast *S. cerevisiae*. In *S. cerevisiae*, the HORMAD protein Hop1 contributes to axial element integrity ^13,14^ and acts in the very early stages of meiotic recombination by mediating the recruitment of the DSB-forming endonuclease Spo11 and its accessory proteins ^15,16^. Hop1 is then phosphorylated by the DNA damage-response kinases Tel1 and Mec1 (homologs of mammalian ATM and ATR, respectively) to recruit the kinase Mek1 ^17^, which enforces DSB repair via the homologous chromosome rather than the sister chromatid ^18,19^. Once the synaptonemal complex assembles, the AAA+ ATPase Pch2 removes Hop1 from the axis ^20–25^, thereby reducing DSB activity and accelerating DSB repair to ultimately enable exit from meiotic prophase and entry into the first meiotic division.

The probability of DSB and CO formation on a given chromosome or chromosomal region varies dramatically across the genome. In some organisms, specific pathways promote axis assembly and/or DSB formation on chromosomal regions at high risk for mis-segregation. In vertebrates, DSB formation at the pseudoautosomal region (PAR) of the X and Y chromosomes is promoted by the vertebrate-specific protein ANKRD31, which directly recruits components of the DSB machinery ^26–28^. Moreover, in *S. cerevisiae* and mammals, the shortest chromosomes experience higher average rates of recombination ^29–32^. In *S. cerevisiae*, this chromosome-size bias is reflected in increased levels of axis protein deposition and DSBs on the shortest three chromosomes, and is abrogated in *hop1* mutants ^33–37^. In addition, *S. cerevisiae* Hop1 also drives enrichment of axis proteins in nucleosome-dense genomic regions, termed “islands”, that are associated with elevated marks of meiotic recombination ^38^. We recently demonstrated that Hop1 enrichment in island regions is driven by an uncharacterized central domain of Hop1 that is predicted to encode a PHD domain ^38^, which in other proteins mediates specific interactions with modified histone tails ^39^.

Here, we determined the structure and function of the Hop1 central domain, which we termed the chromatin binding region (CBR). The Hop1 CBR comprises tightly associated PHD and winged helix-turn-helix (wHTH) domains and directly recognizes the bent DNA on nucleosomes. Mutations that disrupt nucleosome binding cause a loss of Hop1 enrichment from islands, as well as broad defects in DSB formation, which we attribute to the deregulation of distinct DSB-promoting and DSB-inactive conformations of Hop1. Finally, we identify highly divergent CBRs in meiotic HORMADs across eukaryotes, suggesting that these functions may be broadly conserved, with likely functional specialization between eukaryotic lineages.

## Results

### Structure of the budding-yeast Hop1 CBR

To understand the molecular basis for Hop1 CBR-mediated axis enrichment at genomic islands, we sought to determine a high-resolution structure of this domain. We could purify the isolated *S. cerevisiae* Hop1 CBR (residues 322-537) but could not identify crystallization conditions. After screening the equivalent domain from several budding yeast Hop1 proteins, we determined a 1.5 Å-resolution crystal structure of the Hop1 CBR from *Vanderwaltozyma polyspora* (residues 317-535; 34% identical to *S. cerevisiae* Hop1 in this region) (**Table S1**). The *V. polyspora* Hop1 CBR structure reveals a compact assembly with a PHD domain (residues 319-374) and a variant winged helix-turn-helix (wHTH) domain (residues 374-439) with a C-terminal extension that we term HTH-C (residues 440-524; **Figure 1A-B**). The N-terminal PHD domain coordinates two zinc ions through seven conserved cysteine residues and one conserved histidine residue (**Figure 1b**) ^14^. The wHTH and HTH-C domains fold together, with the HTH-C region forming an elaborated β-sheet wing on the wHTH domain and a long C-terminal α -helix stretching across both the PHD and wHTH domains (**Figure 1B**).

**Figure 1.**
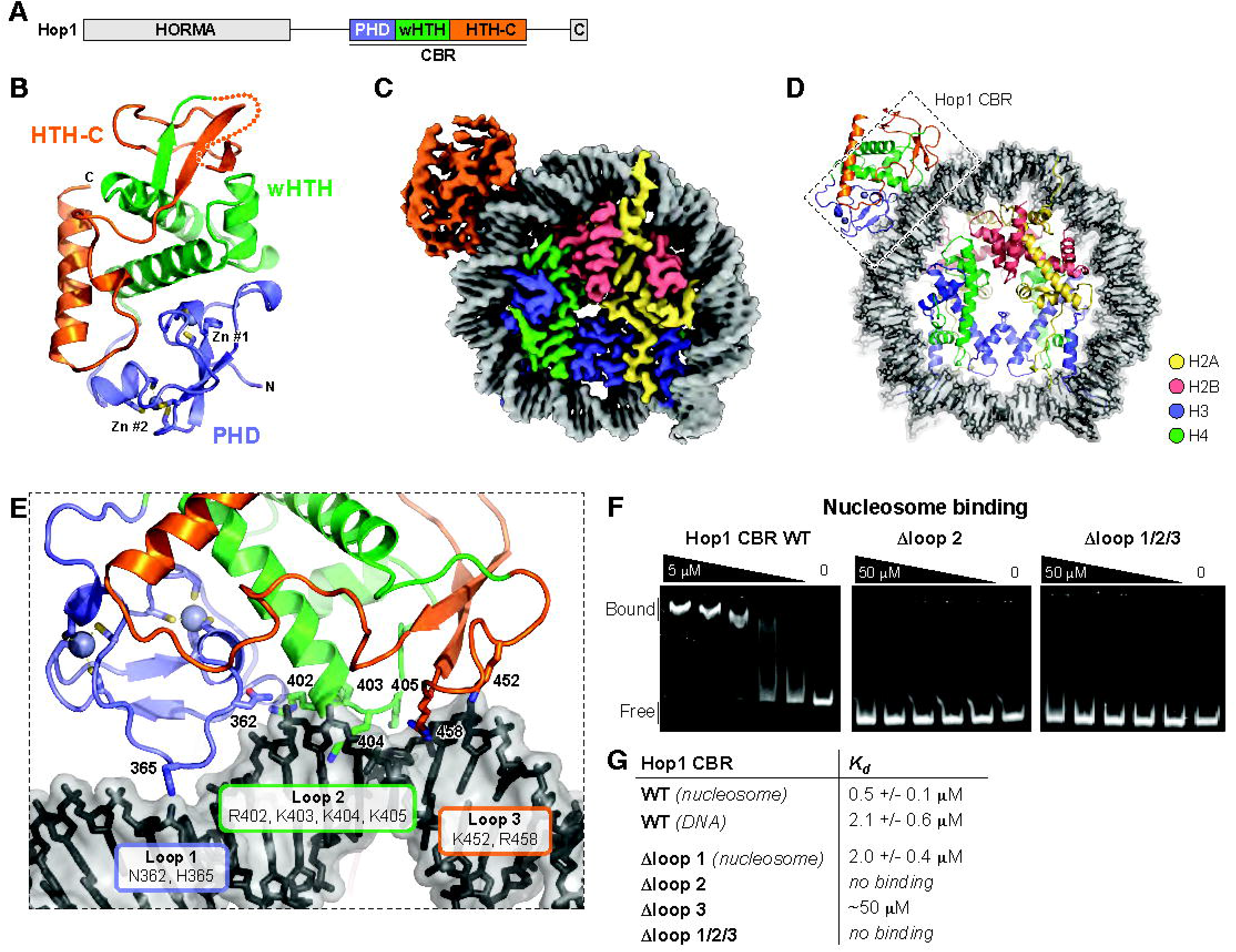
Structure and nucleosome binding by the Hop1 CBR. (A) Domain structure of budding-yeast Hop1. C: closure motif. (B) Crystal structure of the *V. polyspora* Hop1 CBR, with PHD domain colored blue, wHTH green, and HTH-C orange. See **Figure S1** for sequence alignments. (C) CryoEM density map at 2.74 Å resolution of *S. cerevisiae* Hop1 CBR bound to a mononucleosome. (D) Cartoon view of *S. cerevisiae* Hop1 CBR bound to a mononucleosome, oriented as in panel (c). Hop1 is colored as in panel (b), DNA is colored gray/black, and histones are colored yellow (H2A)/red (H2B)/blue (H3)/green (H4). See **Figure S2** for reconstitution of the Hop1 CBR:nucleosome complex, **Figure S3** for cryoEM structure determination workflow, **Figure S4** for structure of a nucleosome with two Hop1 CBRs bound, and **Figure S6** for comparison of *V. polyspora* and *S. cerevisiae* Hop1 CBR structures. (E) Closeup view of the Hop1 CBR-DNA interface, with key DNA-interacting residues of loops 1 (blue), (green), and 3 (orange) shown as sticks and labeled. See **Figure S5** for cryoEM density. (F) Representative electrophoretic mobility shift assays (EMSAs) showing binding of wild-type, Δloop2, and Δloop1+2+3 mutants of *S. cerevisiae* Hop1 CBR to reconstituted nucleosomes. (G) Hop1 CBR-nucleosome and DNA binding affinities determined from triplicate EMSA assays (see **Figure S7** for gels).

PHD domains are found in many families of chromatin-associated proteins, and typically bind modified lysine residues in histone tails via a conserved pocket lined with hydrophobic residues ^39^. Examining our structure of the *V. polyspora* Hop1 CBR reveals that in Saccharomycetaceae Hop1 proteins, the PHD domain’s canonical lysine binding pocket is poorly conserved and largely lacking hydrophobic residues (**Figure S1, Table S2**). This lack of conservation casts doubt on whether the Hop1 PHD domain could bind a histone tail equivalently to other PHD domain proteins. Moreover, comparison of the Hop1 CBR structure with those of DNA-bound wHTH domains reveals that in Hop1, the canonical wHTH DNA-binding surface is partially occluded by the PHD and HTH-C domains (not shown). Thus, if the Hop1 CBR directly binds nucleosome-rich chromatin as suggested by prior studies, it likely does so in a manner unlike canonical PHD or wHTH domain proteins.

### The Hop1 CBR binds nucleosomes

We next directly tested whether the Hop1 CBR binds to nucleosomes *in vitro*. Using a pulldown assay, we detected strong binding of the isolated *S. cerevisiae* Hop1 CBR to reconstituted mononucleosomes (**Figure S2A**). To determine the structural basis for this interaction, we reconstituted a Hop1 CBR:mononucleosome complex (**Figure S2B**) and collected a single-particle cryoelectron microscopy (cryoEM) dataset (**Figure S3, Table S3**). Initial 2D class averages clearly showed nucleosomes, with some classes showing weak density extending outward from the DNA gyres (**Figure S3**). 3D reconstructions revealed two major particle classes corresponding to either a nucleosome or a Hop1 CBR:nucleosome complex. We refined the isolated nucleosome structure to a resolution of 2.57 Å, and the Hop1 CBR:nucleosome complex structure to 2.74 Å (**Figure S3, Table S3**), enabling us to confidently build and refine complete atomic models and visualize the molecular details of the CBR-nucleosome interaction (**Figure 1C-D**). Finally, we detected a minor population with two copies of the Hop1 CBR bound to neighboring locations on a single nucleosome. The low particle number of this class (25,000 particles, compared to 140,000 for the Hop1 CBR:nucleosome complex) precluded full refinement and model building, but we could place two Hop1 CBRs into gaussian-smoothed maps at 3.15 Å resolution to understand the binding of both Hop1 CBR protomers to the nucleosome (**Figure S4**).

Nucleosomes contain two copies of each of four histone proteins – H2A, H2B, H3, and H4 – with 146 bp of DNA wrapped ∼1.75 times around the histone octamer core ^40^. While the histone octamer is two-fold symmetric, the “Widom 601” DNA sequence we used to reconstitute mononucleosomes is asymmetric ^41^. To determine the most likely orientation of this DNA sequence in our structure, we built and refined both possible orientations and chose the one with the higher overall Pearson’s correlation coefficient between the final model and the experimental map. Within the nucleosome structure, locations along each 72-73 base stretch of DNA extending from the midpoint of the 146 bp DNA (the “dyad axis”) are defined as superhelical locations (SHL) 0 at the dyad axis to +/- 7 near the DNA ends ^40^. In our structure, the major Hop1 CBR binding site is between SHL 2.5 and 3.5, on the outer edge of the nucleosome-wrapped DNA (**Figure 1E**). The minor CBR binding site is located between SHL −5.5 and −6.5. The overall structure of the nucleosome-bound *S. cerevisiae* Hop1 CBR closely matches our crystal structure of the *V. polyspora* Hop1 CBR (**Figure S5**), with the most significant difference being that a disordered loop in the HTH-C domain of the *V. polyspora* Hop1 CBR is well-ordered and interacting with DNA in the nucleosome-bound *S. cerevisiae* Hop1 CBR (**Figure 1E**).

The *S. cerevisiae* Hop1 CBR binds DNA through three loops, termed loops 1-3 (**Figure 1E, S6**). Loop 1 is positioned in the PHD domain, with the side-chains of N362 and H365 interacting with the major groove of base pairs 25-27 (as measured from the nucleosome dyad axis; all DNA base pairs noted are for the major Hop1 binding site spanning SHL 2.5-3.5). Loop 2 is in the wHTH domain, with R402, K403, K404, and K405 inserting into the minor groove of base pairs 28-31. Finally, loop 3 is in the HTH-C domain, with K452 and R458 binding base pairs 32-34. Overall, loops 1-3 define a ∼10 bp footprint for the Hop1 CBR on the outer edge of the bent nucleosomal DNA. Notably, despite sharing a common fold with canonical DNA-binding wHTH domains, the Hop1 wHTH domain binds DNA on a distinct surface compared with these domains. Moreover, we do not detect any density for a histone tail bound to the Hop1 PHD domain. Together, these findings point to a model in which the Hop1 CBR recognizes nucleosomes solely through a non-canonical DNA binding surface on its PHD and wHTH domains. While our identification of two distinct binding sites for the Hop1 CBR on the Widom 601 sequence suggests a degree of sequence specificity, we do not observe any sequence-specific interactions in our structure, and can detect no sequence similarity between the two sites (**Figure S4**). These data suggest that binding of the Hop1 CBR to nucleosomes may be driven mainly by recognition of the distinctive bent conformation of DNA in this complex.

We used electrophoretic mobility shift assays (EMSAs) to quantitatively compare binding of the Hop1 CBR to nucleosomes versus a 40-bp DNA segment encompassing the major Hop1 binding site on the nucleosome. The isolated *S. cerevisiae* Hop1 CBR bound nucleosomes with a *K*_*d*_ of 760 nM, compared to 2.1 μM for DNA alone, supporting the idea that the Hop1 CBR specifically recognizes the bent conformation of nucleosomal DNA (**Figure 1F-G, S7**). To test the role of loops 1-3 in nucleosome binding, we generated alanine mutants of the DNA-interacting residues in each loop, plus constructs with two or all three loops mutated. Mutation of loop 1 (N362A/H365A) had the smallest effect on nucleosome binding, reducing the *K*_*d*_ from 760 nM to 2.0 μM (**Figure 1G, S7B**). Mutation of loop 3 (K452A/R458A) had a stronger effect, reducing the *K*_*d*_ to ∼50 μM, and mutation of Loop 2 (R402A/K403A/K404A/K405A) had the strongest effect. with no detectable binding at the highest protein concentration tested (50 μM) (**Figure 1G, S7C-D**). We also detected no binding when combining the loop 2 and 3 mutants, or when combining the loop 1, 2, and 3 mutants (**Figure S7E-H**). Thus, loop 2 is the most important determinant for nucleosome binding by the Hop1 CBR, with loop 3 making a less significant contribution and loop 1 playing only a minor role.

### Nucleosome binding by the CBR directs Hop1 to nucleosome-enriched islands and short chromosomes

The Hop1 CBR specifically promotes axis protein enrichment in nucleosome-dense islands across the yeast genome ^38^. To determine if this enrichment is a consequence of CBR-nucleosome binding, we mutated loop 2 in the endogenous *HOP1* locus. Hop1-loop2 protein (R402A/K403A/K404A/K405A) was expressed at levels and kinetics similar to wild type, indicating that the *hop1-loop2* mutation does not alter *HOP1* expression or meiotic entry (**Figure S8**). In addition, Hop1-loop2 protein exhibited phosphorylation-dependent mobility shifts similar to wild type, indicating that CBR-nucleosome binding is not required for Hop1 phosphorylation ^17^. However, spore viability of *hop1-loop2* cells was reduced (74.8%) compared to wild type (96.1%) indicating that the CBR is important for Hop1 function (**Figure 2A**).

**Figure 2.**
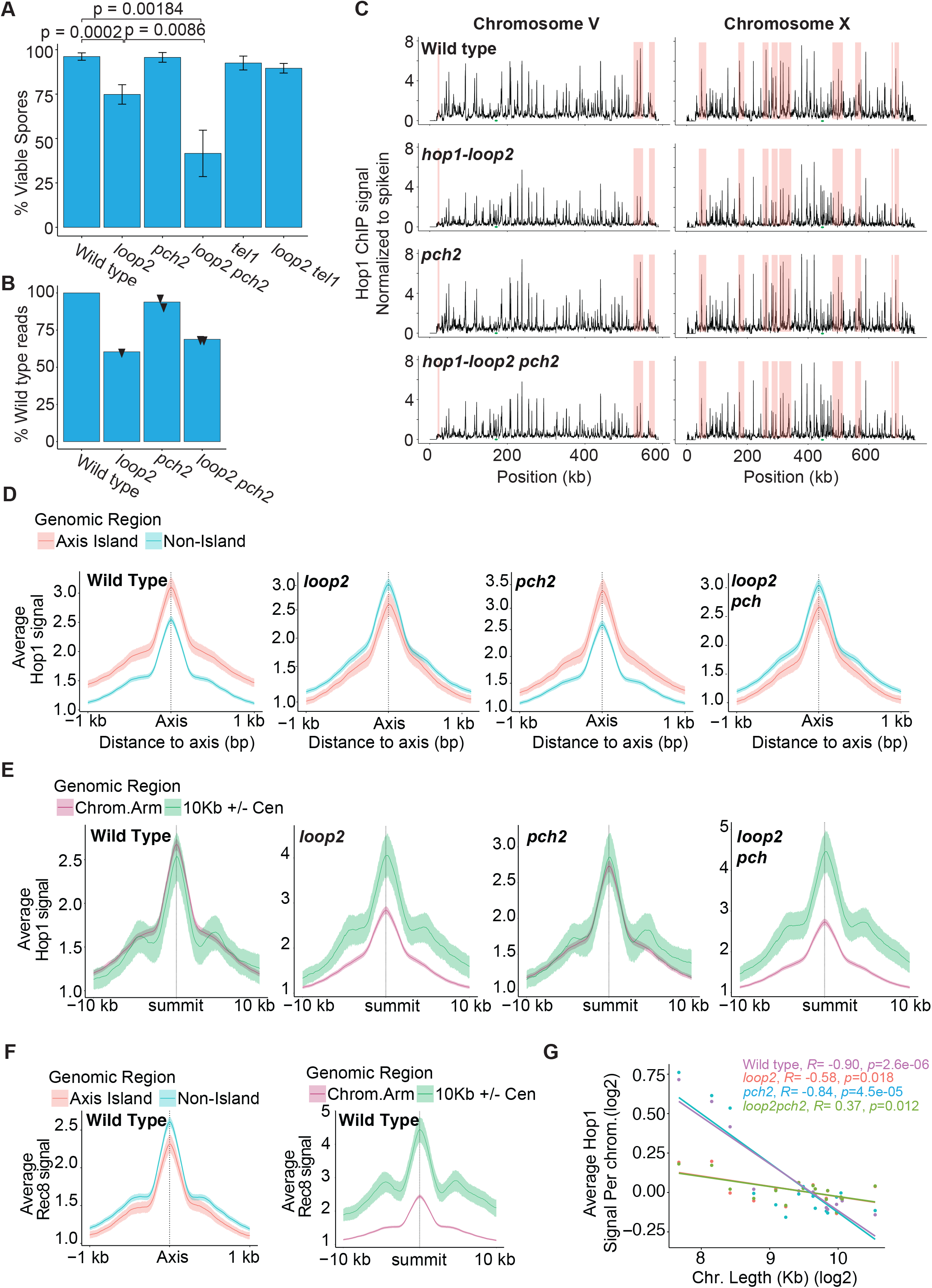
Hop1 association to DNA is altered in the absence of nucleosome binding. (A) Spore viability for each genotype. P values represent the Bonferroni-corrected results of unpaired t-test between groups indicated by connected lines. (B) Read counts relative to wild type, normalized by SNP-ChIP. (C) SNP-ChIP normalized Hop1 signal plotted along chromosome V and chromosome X for each genotype. The centromere on each plot is indicated by a green dot. Island regions are shaded in red. (D)Metaplot analysis of Hop1 ChIP signal at axis attachment sites ^36^ in island or non-island genomic regions. (E) Metaplot analysis of Hop1 ChIP signal at axis attachment sites within 10 kb of the centromere compared to Hop1 ChIP signal at axis attachment sites on chromosome arms. (F) (left) Metaplot of average Rec8 ChIP signal at axis attachment sites in islands vs non-islands in wild type. (right) Metaplot of average Rec8 ChIP signal at axis attachment sites in a wild type within 10 kb of the centromere compared to chromosome arms. (G) Mean Hop1 ChIP-seq enrichment per kb is plotted for each chromosome on a log scale with regression analysis for each genotype.

Analysis of Hop1 localization by spike-in normalized chromatin immunoprecipitation and sequencing (SNP-ChIP normalized ChIP-seq ^42^) revealed that total Hop1 chromatin association in *hop1-loop2* cells was reduced to 60% of wild type, with a specific loss of Hop1 enrichment from island regions (**Figure 2B-C**). In wildtype, the Hop1 binding in island regions is 1.56 fold higher than Hop1 binding in desert regions. In *hop1-loop2* cells, the Hop1-loop2 binding in islands is about the same as Hop1-loop2 binding in desert regions (∼1.04 fold). These results demonstrate that the enrichment of Hop1 in nucleosome-dense regions of the yeast genome ^38^ is a direct consequence of nucleosome binding by Hop1.

In addition to the overall reduction of Hop1-loop2 on chromosomes, we observed a specific enrichment of Hop1-loop2 around centromeres compared to wild-type Hop1 (**Figure 2C, E**). Meta-analysis of axis association sites in centromere regions, defined here as the centromere +/- 10 kb, showed higher Hop1-loop2 association compared to chromosome arm regions (**Figure 2E**). This enrichment likely reflects axis-protein recruitment by Rec8-containing cohesin complexes, which are similarly enriched around centromeres in wild type cells (**Figure 2F**) and represent the only remaining axis-protein recruitment pathway in mutants lacking Hop1 CBR function ^38^.

In addition to island regions, Hop1 also mediates increased association of axis protein on the three shortest *S. cerevisiae* chromosomes ^36^. To test whether the Hop1 CBR plays a direct role in this chromosome-size bias, we determined the relative Hop1 enrichment in *hop1-loop2* mutants as a function of chromosome size. In wild type cells, Hop1 was enriched on the shortest three chromosomes, but this effect was strongly weakened in *hop1-loop2* mutants (**Figure 2G**). These results demonstrate that Hop1 CBR-nucleosome binding drives key aspects of the chromosome distribution of Hop1, including enrichment on axis islands and on short chromosomes.

### Mutation of the Hop1 CBR leads to defective homolog synapsis

To further assess Hop1 localization on meiotic chromosomes, we prepared chromosome spreads from meiotic prophase cells (2, 3, and 4 hours after meiotic induction) and probed them with antibodies against Hop1 and the SC central-region protein Zip1, a marker of chromosome synapsis. Compared to wild type, *hop1-loop2* nuclei in early prophase (hour 2) only contained about 26.9% of Hop1 signal intensity (**Figure 3A-B**), and the nuclear area Hop1 occupied (measured as a percentage of total DAPI area) in *hop1-loop2* cells was only about 20% of the nuclear area occupied by wild type Hop1 (**Figure 3A, C**), indicating a defect in the chromosomal association of Hop1. These results do not reflect differences in synchrony because meiosis initiated at the same time in both strains (**Figure S8B**).

**Figure 3.**
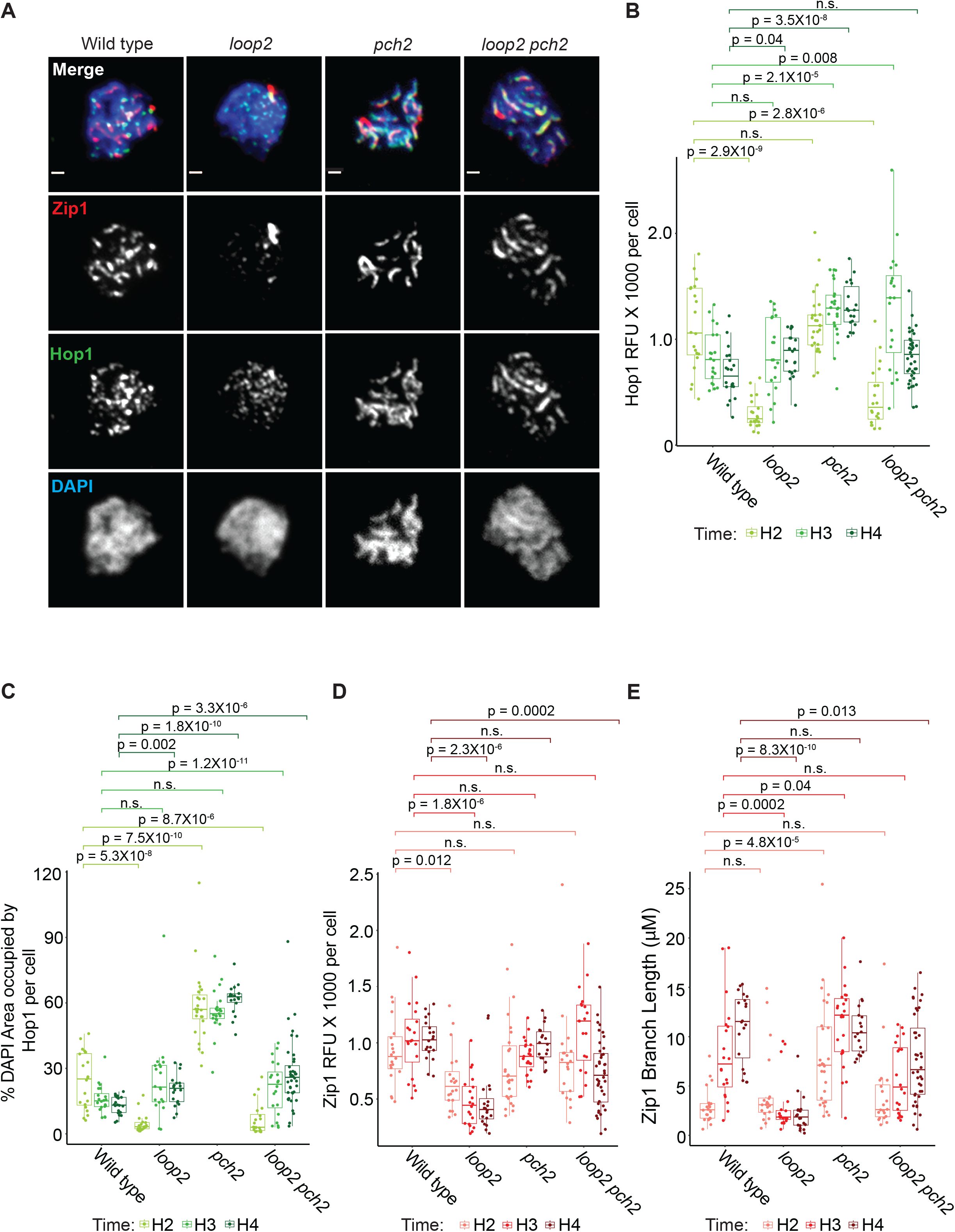
Hop1 CBR domain is important for synapsis. (A) Representative images from chromosome spreads prepared at 3 hours after meiotic induction, hybridized with antibodies against Zip1 and Hop1, stained with DAPI. Scale bars: 1 μm. (B) Dot plot of the quantification of total Hop1 relative fluorescent units (RFU) per cell. Hours 2, 3, and 4 are separated by color as depicted in the legend below. Standard deviation and means are displayed for each as a box plot. Lines between categories represent p value results; n.s. is not significant. Statistics are determined by unpaired Wilcoxon test with Bonferroni correction. (C) Dot plot of the % total DAPI area that the Hop1 signal occupies for each cell. Plot layout and statistics are the same as panel (B). (D) Dot plot of the quantification of total Zip1 RFU per cell. Plot layout and statistics are the same as panel (B). (E) Zip1 length for each cell is displayed in micrometers. Plot layout and statistics are the same as panel (B).

By late prophase, chromosomal Hop1 levels in wild-type cells declined as chromosomes as a consequence of chromosome synapsis ^23^. By contrast, *hop1-loop2* mutants failed to undergo proper chromosome synapsis and accumulated chromosomal Hop1 similar to other mutants with synapsis defects (**Figure 3A-C**) ^23^. To quantify the SC defect of *hop1-loop2* mutants, we classified nuclei by their Zip1 structures. Whereas the majority of wild-type nuclei displayed full Zip1 polymerization on chromosomes at hour 3 and 4 of meiosis, *hop1-loop2* nuclei displayed primarily punctate Zip1 signal or short Zip1 tracks, similar to Zip1 patterns seen at hour 2 in wild type, and often contained one or two Zip1 aggregates (polycomplexes; **Figure 3A,D-E**). Thus, *hop1-loop2* cells fail to form wild-type SC structures.

### *hop1-loop2* cells experience decreased DSB formation

To investigate the functional consequences of altered axis assembly and dynamics in *hop1-loop2* mutants, we assayed chromosome-wide DSB induction by pulsed-field gel electrophoresis and Southern blotting. To exclude differences in DSB turnover, we performed this experiment in a *rad50S* background, which blocks DSB repair ^43^. Quantification of DSB signals along chromosome 8 revealed an approximately 50% reduction in DSB formation in *hop1-loop2* mutants compared to wild-type (**Figure 4A**). This reduction was less severe than seen in the complete absence of *HOP1*, indicating that Hop1 is able to promote some DSB formation even in the absence of nucleosome binding. We observed a similar 30-50% reduction in DSB levels in the *hop1-loop2* mutant when measuring chromosomal fragments below the smallest chromosome as a proxy for whole-genome DSB formation (**Figure 4B**), and when measuring DSB levels at the engineered *HIS4:LEU2* hotspot in *RAD50* cells (**Figure 4C**). However, not all DSB hotspots were equally affected, as DSB levels were nearly unaffected at the *GAT1* hotspot on chromosome 6 (**Figure 4D**). These data indicate that Hop1 nucleosome binding promotes DSB formation in some, but not all, regions of the genome.

**Figure 4.**
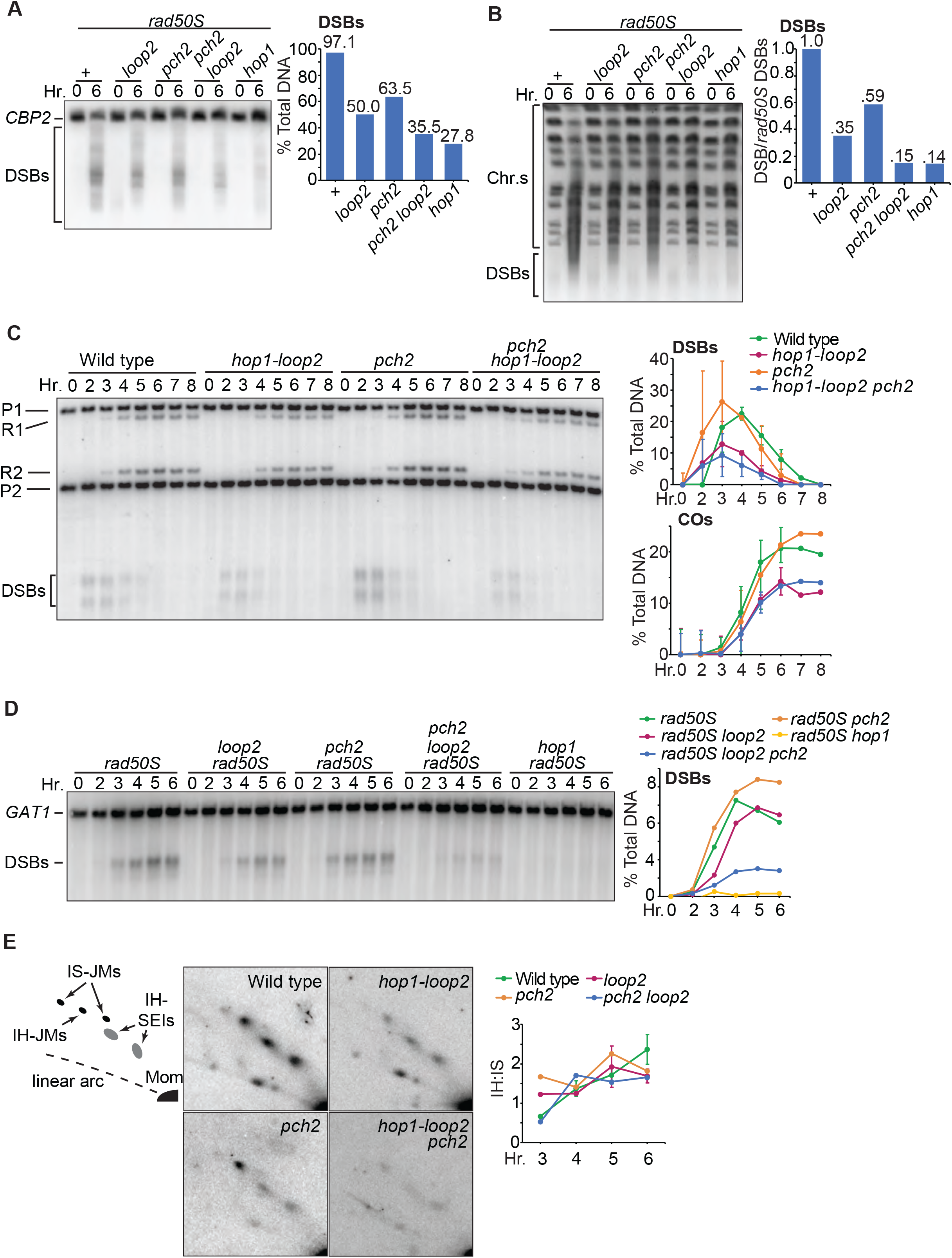
Hop1 CBR domain promotes DSB activity, but not repair choice. (A) (Left) Pulsed-field gel and Southern blot analysis of chromosome VIII (analyzed using a probe against *CBP2*). (Right) Percent total broken DNA calculated based on a Poisson distribution using the intact chromosome VIII bands for each genotype and assuming full replication by 6 hours ^78^. (B) (Left) Pulsed-field gel resolving all chromosomes for each genotype stained with ethidium bromide. (Right) DSB levels measured below the smallest chromosome band are compared between genotypes to estimate total genomic DSB levels. (C) (Left) Southern blot analysis of the *HIS4:LEU2* hotspot locus across genotypes during meiosis. P1 and P2 represent parental alleles. R1 and R2 represent COs. DSBs appear near the bottom of the gel. (Right Top) Quantification of DSBs at the *HIS4:LEU2* locus shown as a percentage of total DNA signal for each lane. (Right Bottom) Measurements of recombination between P1 and P2 depicted as a percentage of total DNA. (C) (Left) Southern blot analysis of the *GAT1* locus across genotypes during meiosis. Top band represents the intact *GAT1* locus. Bottom band represents DSBs that occur at the *GAT1* locus. (Right) Measurements of the DSB induction for each genotype at the *GAT1* locus. (E) Native-native 2D gel electrophoresis and Southern blot analysis of psoralen-stabilized recombination intermediates at the *HIS4:LEU2* locus. (Left) Illustration of the expected pattern of recombination intermediates. IS-JM - intersister join molecules, IH-JM - interhomolog joint molecules. SEIs - single end invasion intermediates, a precursor of IH-JMs. (Middle) representative image for each genotype at hour 4. (Right) Measurements of joint molecule signals depicted as the ratio of IH-JM signal to IS-JM signal.

### Nucleosome binding by Hop1 is not necessary for DSB repair template bias

In *hop1*Δ mutants, the few DSBs that do form are repaired via the sister chromatid, instead of the homologous chromosome, resulting in the loss of COs ^44^. To investigate the function of CBR-mediated Hop1 recruitment during meiotic DSB repair, we analyzed meiotic recombination at *HIS4-LEU2* ^45^. Southern blot analysis of the *hop1-loop2* mutant revealed that CO levels were reduced approximately 50% compared to wild type, mirroring the reduction in DSB levels at this locus (**Figure 4C**). To test if CO repair is normal in the absence of Hop1-nucleosome binding, we measured stabilized recombination intermediates using 2D gel electrophoresis and Southern blotting. The ratio of joint-molecule repair intermediates between homologous chromosomes and sister chromatids in *hop1-loop2* mutants was comparable to wild-type cells (**Figure 4E**). Thus, repair template choice, another key event regulated by Hop1, does not require nucleosome binding by the CBR, functionally separating the CBR from the neighboring SQ/TQ cluster domain (SCD; residues 298-318), which is essential for repair template choice ^17^.

### Pch2 depletes Hop1-loop2 from meiotic chromosomes

The reduced chromosomal association of Hop1 seen in *hop1-loop2* mutants raises the possibility that Hop1-loop2 protein is a better target for removal by the Hop1 disassemblase Pch2. Indeed, we observed increased levels of Hop1 on chromosome spreads in *hop1-loop2 pch2* double mutants (**Figure 3A-C**). The return of Hop1 levels in the *hop1-loop2 pch2* double mutant is likely attributable tothe absence of Pch2 disassembly activity as well as the elevated Hop1 protein levels seen in *pch2* mutants (**Figure S8A**) ^46^. Notably, deletion of *PCH2* in *hop1-loop2* mutants did not lead to greater ChIP signal on chromosomes and did not rescue Hop1 binding to island regions (**Figure 2B-D**), indicating that deletion of *PCH2* did not restore the ability of Hop1 to directly bind to chromatin.

In line with the recovery of Hop1 binding to meiotic chromosome spreads, deletion of *PCH2* also nearly restored SC formation in *hop1-loop2* mutants (**Figure 3A**). Total Zip1 signal intensity and Zip1 tract length in *hop1-loop2 pch2* cells was comparable to wildtype in hour 2 and hour 3 after meiotic induction (**Figure 3A,D-E**), and chromosomes also showed nearly wild-type localization of the the SC central-element protein Gmc2 ^47^, indicating that *hop1-loop2 pch2* double mutants form bona fide SCs (**Figure S9**). Both Zip1 signal intensity and Zip1 length were lower than wild-type Zip1 levels at hour 4, suggesting that these cells may not achieve the same extent of synapsis as observed in wild type (**Figure 3A,D-E**). Nonetheless, these results indicate that nucleosome binding by Hop1 is not necessary for meiotic chromosome synapsis. Rather, the defect in SC formation in *hop1-loop2* cells is at least in part a consequence of premature Hop1 removal by Pch2.

### *hop1-loop2* spore survival and DSB formation depends on Pch2

If the *hop1-loop2* phenotypes were solely due to increased Pch2-mediated Hop1 removal from chromosomes, then deletion of *PCH2* should also rescue the DSB defect of the *hop1-loop2* mutant. Contrary to this expectation, we found that DSB levels were further reduced in *hop1-loop2 pch2* double mutants (**Figure 4A-D**). The DSB reduction was observed both by pulsed-field gel electrophoresis and by analysis of individual loci, although we again observed differences in effect size at individual hotspots. *hop1-loop2 pch2* mutants showed only mildly reduced the DSB levels compared to *hop1-loop2* single mutants at *HIS4-LEU2*, but severely reduced DSB levels at *GAT1*, suggesting locus-specific differences in each hotspot’s dependence on Hop1 CBR-nucleosome binding and Pch2. Of the DSBs that did occur at *HIS4-LEU2* in the *hop1-loop2 pch2* background, the bias toward inter-homolog repair remained intact (**Figure 4E**). The drop in DSB formation was associated with a major decrease in spore survival in the double mutant background (wild type viability = 96.1%, *hop1-loop2 pch2* viability = 41.6%; *P* = 0.00183, unpaired t-test) (**Figure 2B**). These results indicate that *hop1-loop2* mutants depend on *PCH2* to support DSB formation. Notably, *TEL1* deletion, which increases the levels of meiotic DSB formation ^48,49^, was sufficient to restore the spore viability of *hop1-loop2* mutants to near wild-type levels (**Figure 2A**). The simplest explanation for these observations is that Hop1 exists in two distinct conformational states that differ in their ability to promote DSB formation and whose relative prevalence is regulated by nucleosome binding and Pch2-dependent remodeling (see **Discussion**).

### Meiotic HORMADs across eukaryotes possess diverse CBRs

Our data show that the *S. cerevisiae* Hop1 CBR plays important roles in recombination, but this domain is notably lacking in meiotic HORMADs from other model organisms like *C. elegans* and *M. musculus*. To determine how widespread the CBR is among meiotic HORMADs, we generated profile hidden Markov Models (HMMs) for the PHD, wHTH, and HTH-C domains in *S. cerevisiae* Hop1, then searched for similar domains in meiotic HORMADs from a set of 158 diverse eukaryotes ^50,51^. We selected 147 meiotic HORMADs from 158 genomes and transcriptomes representative of the full diversity of the eukaryotic tree of life (thirteen species encode two meiotic HORMADs, and *C. elegans* encodes four). Those genomes that lack an identifiable meiotic HORMAD protein also usually lack homologs of the chromosome axis core proteins (e.g. *S. cerevisiae* Red1 or *M. musculus* SYCP2/SCYP3; ^52^). The meiotic HORMADs we could identify almost always encode both an N-terminal HORMA domain and a C-terminal closure motif, but show a strikingly variable architecture within their central regions as detected by HMM profile matches and AlphaFold structure predictions (**Figure 5A-B, TableS4**) ^53,54^. The majority of HORMADs we identified (106 of 147) encode a CBR some combination of a PHD and a wHTH domain. Across Saccharomycetaceae – the fungal group that contains *S. cerevisiae* – we detected a conserved CBR architecture with PHD, wHTH, and HTH-C domains. In a large set (36/147 of meiotic HORMADs from Opisthokonta (which includes Fungi and Metazoa), the CBR comprises predicted PHD and wHTH domains, but lacks HTH-C (**Figure 5A-B, Figure S10**). In these proteins, the PHD domain’s canonical histone tail binding hydrophobic cage is highly conserved (**Figure S10A-B**), and the wHTH domain’s canonical DNA binding site is predicted to be surface-exposed and positively charged (**Figure S10C-E**). These data suggest that, unlike *S. cerevisiae* Hop1, the larger group of CBRs lacking HTH-C may bind nucleosomes in a bipartite manner through both histone tails and DNA. In the pathogenic fungal species *Encephalitozoon cuniculi* and *Encephalitozoon intestinalis*, we identified putative meiotic HORMADs that lack an N-terminal HORMA domain, and encode only a CBR with a PHD/wHTH domain or PHD domain, respectively (**Figure 5A-B, Table S4**).

**Figure 5.**
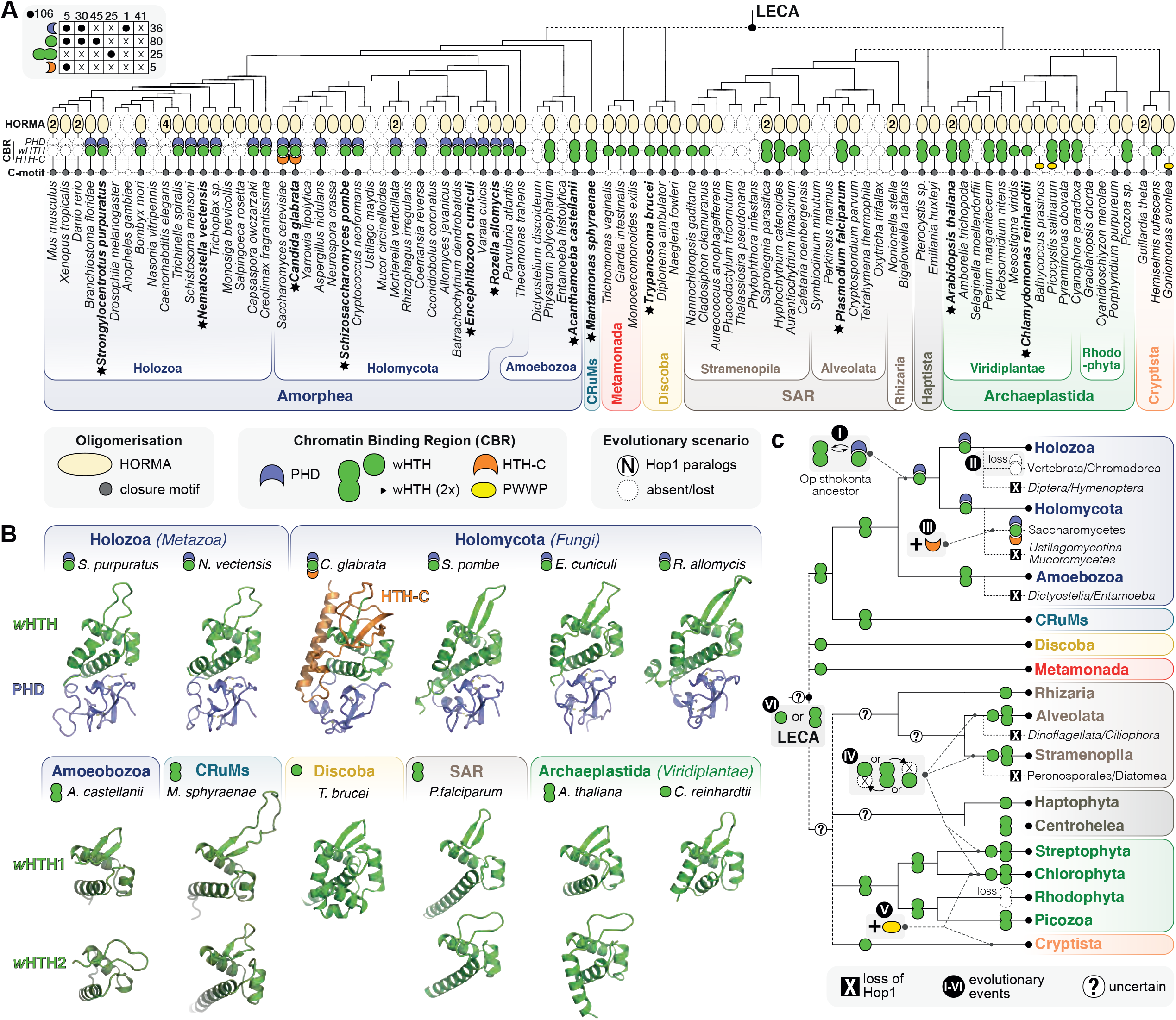
Meiotic HORMADs encode a variable central chromatin-binding region. (A) Cartoon of a phylogenetic tree of diverse eukaryotes (a subset of 87 out of 158 species analyzed in this study), with individual domains in meiotic HORMAD proteins noted: HORMA domain beige, PHD domain blue, wHTH domain green, HTH-C domain orange, PWWP yellow and the closure motif grey. Numbers in the HORMA domain field indicate the number of detected meiotic HORMADs in a given genome. Two different shapes are depicted for the wHTH domain, each indicating the number of wHTH domains detected in a single meiotic HORMAD protein. Major eukaryotic groups are color-coded and labeled. Top left is a presence-absence table for the key domains of the Chromatin-Binding Region (CBR) of all 147 HORMAD homologs analyzed in this study. See full dataset of 158 species/147 HORMAD proteins in **Table S4**. (B) AlphaFold 2 predicted structures of selected meiotic HORMAD CBRs (indicated with a star and in bold text in panel (A)). All CBRs are shown with PHD domain blue, wHTH green, and HTH-C orange. *Strongylocentrotus purpuratus* (Uniprot ID A0A7M7T0L9) residues 329-476 (PHD 329-395, wHTH 396-476); *Nematostella vectensis* (Uniprot ID A7RLI6) residues 280-426 (PHD 280-345, wHTH 346-426); *Candida glabrata* (Uniprot ID Q6FIP6) residues 319-526 (PHD 319-373, wHTH 374-438, HTH-C 439-526); *Schizosaccharomyces pombe* (Uniprot ID Q9P7P2) residues 333-480 (PHD 333-388, wHTH 389:480); *Encephalitozoon cuniculi* (Uniprot ID Q8SWC8) resides 1-142 (full length; PHD 1-61, wHTH 62-142); *Rozella allomycis* (Uniprot ID A0A075AU51) residues 399-551 (PHD 399-454, wHTH 455-551); *Acanthamoeba castellanii* (Uniprot ID L8H1D9) residues 246-309 (wHTH 1) and 335-394 (wHTH 2); *Mantamonas sphyraenae* (ID see **Table S4**) residues 254--340 (wHTH 1) and 3787-465 (wHTH 2); *Trypanosome brucei* (Uniprot ID Q38B55) residues 292-406 (wHTH); *Plasmodium falciparum* (Uniprot ID Q8IEM0) residues 973-1053 (wHTH 1) and 1214-1287 (wHTH 2); *A. thaliana* ASY1 (Uniprot ID F4HRV8) residues 297-362 (wHTH 1) and 388-471 (wHTH 2); *Chlamydomonas reinhardtii* (Uniprot ID A0A2K3DBA0) residues 443-516 (wHTH). Of note: *C. reinhardtii* wHTH is most similar to *A. thaliana* wHTH 1, while *T. brucei* wHTH is more similar to other wHTH 2 domains. See **Figures S10** for sequence alignments. (C) Construction of the evolution of the CBR of meiotic HORMAD protein throughout the eukaryotic tree of life. Six key events can be discerned: [I] replacement of the first wHTH domain in the ancestor of fungi and animals (Opisthokonta) with a PHD domain; [II] loss of the PHD-wHTH configuration in a number of animal lineages (i.e. Vertebrata and a Nematoda class Chromadorea); [III] C-terminal extension of the wHTH specific for *Saccharomycetes*; [IV] differential loss of one of the two wHTH domains in various clades within the supergroups SAR and Archaeplastida; [V] independent gains of PWWP-like domains in Chloroplastida and Cryptista; [VI] unclear (inferred) ancestral presence of a single or double wHTH configuration in the last eukaryotic common ancestor (LECA).

In addition to CBRs encoding PHD and wHTH domains, a large set of meiotic HORMADs encode CBRs with either a single wHTH domain (45/106, scattered throughout the tree)) or a tandem repeat of two such domains (25/106; in *Mantamonas*, some Amoebozoa, SAR, and Archaeplastida) (**Figure 5A-B, Figure S11, Table S4**). Notably, both *Arabidopsis thaliana* ASY1 and *Oryza sativa* PAIR2 encode two wHTH domains. These domains may bind DNA in a sequence-specific or nonspecific manner to promote meiotic chromosome axis assembly similarly to CBRs with PHD and wHTH domains.

Based on the conserved domains within the meiotic HORMAD central region, we propose that the last common ancestor of eukaryotes encoded a meiotic HORMAD protein with one or two wHTH domains. In Opisthokonta (Holozoa+Holomycota), this architecture was elaborated to include a PHD domain situated N-terminal to a single wHTH domain (wHTH is most similar to the C-terminal wHTH of double wHTH HORMADs), then further elaborated in Saccharomycetaceae to include the HTH-C domain (**Figure 5C**). The addition of the HTH-C domain also apparently coincided with the development of the distinct DNA binding surface we observe by cryoEM. Intriguingly, we also observe yet other chromatin-binding domains, such as a putative methyl-binding PWWP-like domain (Chlorophyta, Cryptista) or unknown domains (*Aurantiochytrium*, **Figure 5C, Table S4**). Many organisms, notably including mammals and most nematodes, have since lost the CBR entirely, and their meiotic HORMADs apparently encode only an N-terminal HORMA domain and a disordered C-terminal region with a closure motif at the protein’s extreme C-terminus. Strikingly, however, the majority of meiotic HORMADs across eukaryotes encode a CBR with predicted DNA and/or chromatin binding activity.

## Discussion

We have shown here that the Hop1 CBR directly associates with nucleosomes *in vitro* and functions *in vivo* to regulate Hop1 chromatin localization as well as Hop1 activities in both recombination and SC formation. For any given organism, the probability for CO repair is not equal across the genome. Some organisms have evolved additional pathways to promote axis assembly and/or DSB formation on particular chromosome regions at high risk for mis-segregation. In *S. cerevisiae*, the CBR-nucleosome association promotes the DSB activity of Hop1, and by doing so, drives the enrichment of recombination in nucleosome dense island regions of the genome. It remains to be seen if increased recombination in islands provides cells with any evolutionary advantage, although the loss of axis protein enrichment on small chromosomes in the absence of CBR-nucleosome binding suggests that this domain could play an important role in ensuring all chromosomes receive at least one CO.

The increased Hop1 binding to islands and short chromosomes is a direct consequence of the CBR-nucleosome interaction. Our structural data indicate that binding occurs primarily through interactions with the DNA backbone, with limited sequence specificity as indicated by the interaction of loop1 with the major groove and the preferential binding to certain positions along the Widom 601 DNA sequence. The notion that Hop1 binds to nucleosomes relatively non-specifically is supported by *in vivo* data, which shows that cohesin-independent Hop1 enrichment correlates well with regional nucleosome density ^38^. Interestingly, nucleosome binding also influences Hop1 enrichment around centromeres. In wild-type cells, Hop1 and its binding partner Red1 are surprisingly not enriched around centromeres, given the large amount of Rec8-cohesin present in these regions ^36^. By contrast, in *hop1-loop2* mutants, Hop1 enrichment at centromeres mirrors that of Rec8. This centromeric enrichment may simply reflect the fact that Rec8-cohesin is the only remaining axis recruitment mechanism in the absence of Hop1 CBR activity ^38^, but it is possible that the CBR-nucleosome interaction specifically disfavors Hop1 enrichment around centromeres. Centromeres were recently implicated as driving the enrichment of axis-dependent recombination activity on short chromosomes ^33^. It remains to be seen whether the loss of Hop1 enrichment on short chromosomes observed in *hop1-loop2* mutants is a reflection of, or involved in, this mechanism.

In addition to affecting Hop1 distribution, nucleosome binding also impacts the ability of Hop1 to regulate DSB formation and synapsis. Unlike Hop1 distribution, these effects of nucleosome binding are strongly and differentially modulated by Pch2. We propose that these behaviors reflect two states of chromosomal Hop1 whose interconversion is regulated by nucleosome binding and Pch2-dependent disassembly (**Figure 6**). One state, which predominates in early prophase and which we refer to as ‘state 1’, promotes DSB formation and may be refractive to Pch2-dependent disassembly. In contrast, ‘state 2’ is unable to promote DSB formation and susceptible to disassembly by Pch2. In this model, the CBR-nucleosome interaction stabilizes Hop1 in state 1 (**Figure 6**). When the CBR domain is unbound, as in the *hop1-loop2* mutant, Hop1 more readily transitions from state 1 to state 2, enabling Pch2-mediated removal from chromatin (**Figure 6**). In *hop1-loop2 pch2* double mutants, Hop1 accumulates in state 2, leading to extensive Zip1 deposition but preventing the recycling of Hop1 to promote further DSB formation. The existence of two chromosomal forms of Hop1 is supported by cytological data from yeast and *Arabidopsis* 55,56. However, how these two forms of Hop1 differ structurally remains to be determined. Hop1 adopts different protein-protein complexes by engaging its C-terminal ‘safety belt’ region with closure motifs in Red1 and possibly other Hop1 monomers to form multi-protein complexes ^57^. A single Hop1 monomer can likely also form a “self-closed” state, with its HORMA domain bound to its own C-terminal closure motif. In addition, Hop1 is extensively modified by phosphorylation and SUMOylation, creating additional forms of Hop1 that may differ in their ability to promote meiotic DSB formation and their susceptibility to Pch2-mediated disassembly 17,58,59.

**Figure 6.**
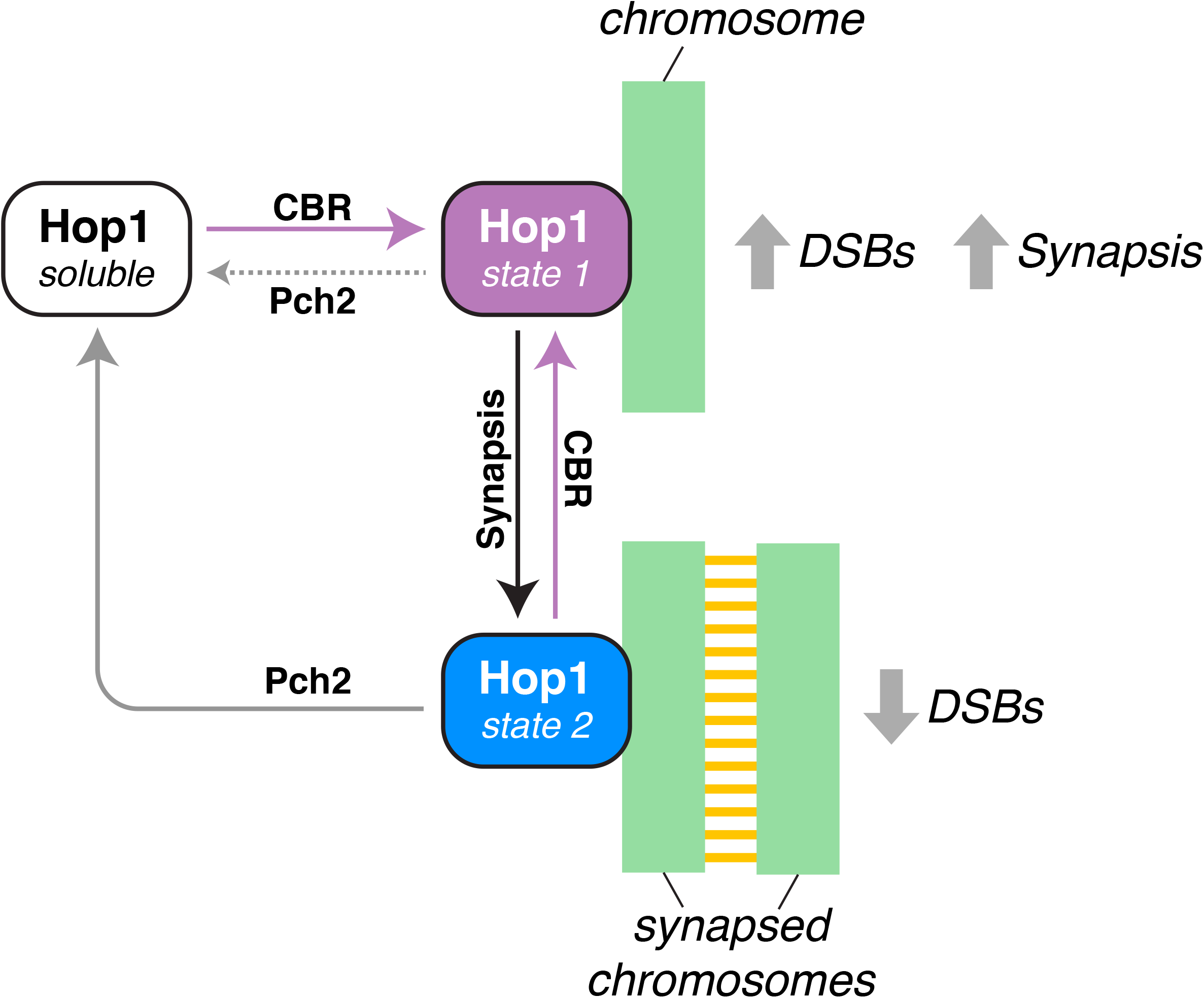
Hop1-CBR nucleosome interaction regulates Hop1 function. We propose that Hop1 exists in at least three states: a soluble state primed by Pch2 for chromatin association (white), and two chromatin-associated states (State 1 purple, State 2 blue). State 1 predominates upon initial chromatin association and promotes DSB formation, which in turn promotes synapsis. State 2, which does not promote DSB formation, predominates after synapsis. Pch2 can readily access and remodel Hop1 in state 2, but state 1 is refractory to Pch2. The Hop1 CBR prevents Hop1 remodeling by stabilizing Hop1 in state 1, preventing the switch to state 2. In the absence of CBR-nucleosome binding, a premature switch from state 1 to state 2 occurs. This switch results in lower DSB levels and premature conversion of Hop1 into the soluble form, destabilizing the SC. Some recombination activity is salvaged by Pch2-mediated recycling Hop1. In the absence of both CBR-nucleosome binding domain and Pch2, Hop1 accumulates in state 2 without recycling, resulting in further decreased DSB formation while permitting SC assembly.

Meiotic HORMADs across eukaryotes possess CBRs with variable architecture: fungi and many animals encode a CBR with tandem PHD and wHTH domains, while most other eukaryotes encode CBRs with one or two wHTH domains. Thus, meiotic HORMADs share a broadly-conserved module capable of direct chromatin binding, which can complement cohesin-mediated axis assembly and plays a key regulatory role in meiotic recombination.

## Materials and Methods

### Protein Expression and Purification

Ligation-independent cloning was used to clone *S. cerevisiae* Hop1^322-537^ and *V. polyspora* Hop1^319-529^ into UC Berkeley Macrolab vectors to generate N-terminal TEV protease-cleavable His6- or His6-maltose binding protein tags (Addgene #29666, 29706). DNA binding mutants of *S. cerevisiae* Hop1^319-529^ (loop 1: N362A+H365A; loop 2: R402A+K403A+K404A+K405A; loop 3: K452A+R458A) were generated by PCR mutagenesis of His6-MBP tagged constructs.

For protein expression, plasmids encoding Hop1 fragments were transformed into *E. coli* LOBSTR cells (Kerafast), and grown in 2XYT media supplemented with 10 μM ZnCl_2_. Cells were grown at 37°C to an OD600 of 0.6, then protein expression induced with 0.25 mM IPTG and cells were grown a further 16 hours at 20°C.

For protein purification, cells were harvested by centrifugation, suspended in resuspension buffer (20 mM Tris-HCl pH 7.5, 300 mM NaCl, 20 mM imidazole, 2 mM β-mercaptoethanol, and 10% glycerol) and lysed by sonication. The lysate was clarified by centrifugation, then the supernatant was loaded onto a Ni2+ affinity column (HisTrap HP, Cytiva) pre-equilibrated with resuspension buffer. The column was washed with resuspension buffer, followed by low-salt wash buffer (20 mM Tris-HCl pH 7.5, 100 mM NaCl, 20 mM imidazole, 2mM β-mercaptoethanol, and 10% glycerol) and eluted with low-salt wash buffer supplemented with 250 mM imidazole. The eluate was loaded onto an anion-exchange column (Hitrap Q HP, Cytiva) and eluted with a gradient to high-salt wash buffer (20 mM Tris-HCl pH 7.5, 1 M NaCl, 20 mM imidazole, 2 mM β-mercaptoethanol, and 10% glycerol). Fractions containing desired protein were pooled and concentrated to 500 μL by ultrafiltration (Amicon Ultra-15, EMD Millipore), then passed over a size exclusion column (HiLoad Superdex 200 PG, Cytiva) in size exclusion buffer (20 mM Tris-HCl pH 7.5, 300 mM NaCl, 10% glycerol, and 1 mM DTT). For crystallization experiments, the N-terminal His6-tag on *V. polyspora* Hop1^317-535^ was cleaved by TEV protease treatment (16 hours at 4°C) of pooled anion-exchange fractions. The cleavage reaction was passed over a second Ni2^+^ affinity column and the flow-through containing cleaved protein was collected, concentrated, and further purified by size exclusion. Purified proteins were concentrated by ultrafiltration and stored at 4°C for crystallization, or aliquoted and frozen at −80°C for biochemical assays. All mutant proteins were purified as wild-type.

### Nucleosome core particle reconstitution

Lyophilized *Xenopus laevis* histone proteins (H2A, H2B, H3, and H4) were purchased from the Histone Source at Colorado State University (https://histonesource-colostate.nbsstore.net). Histone octamer and full nucleosome assembly was performed essentially as described ^60^. Briefly, histone octamers were assembled by resuspending individual purified histones in unfolding buffer (6 M Guanidinium HCl, 20 mM Tris-HCl pH 7.5, 5 mM DTT) to a concentration of 2 mg/mL, then mixing in a molar ratio of 1 H2A:1 H2B:0.9 H3:0.9 H4 and diluted to a final concentration of 1 mg/mL. The mixture was placed in a dialysis cassette (Pierce Slide-A-Lyzer, 3.5 kDa MWCO) and dialyzed three times (4h, 12h, 4h) into refolding buffer (2 M NaCl, 10 mM Tris-HCl pH 7.6, 1 mM EDTA, 1 mM DTT). Assembled octamers were separated from unassembled H2A:H2B dimer by size exclusion (Superdex 200 PG) in refolding buffer.

A 146-bp DNA fragment corresponding to the Widom 601 DNA sequence ^41^ (sequence below) was amplified by PCR and purified by anion exchange chromatography (HiTrap Q HP, Cytiva). DNA and histone octamers were assembled by overnight dialysis from 1.4 M TEK buffer (1.4 M KCl, 10 mM Tris-HCl pH 7.5, 0.1 mM EDTA, 1 mM DTT), to 10 mM TEK buffer (10 mM KCl, 10 mM Tris-HCl pH 7.5, 0.1 mM EDTA, 1 mM DTT). Fully-assembled nucleosomes were purified by size exclusion (Superose 6 10/300 GL, Cytiva) to ensure quality and purity.

>Widom 601 DNA sequence (chain J in models; reverse complement is chain I in models)

TGGAGAATCCCGGTGCCGAGGCCGCTCAATTGGTCGTAGACAGCTCTAGCACCGCTTAAACGCACGTACGCGCTGTCCCCCGCGTTTTAACCGCCAAGGGGATTACTCCCTAGTCTCCAGGCACGTGTCAGATATATACATCCTGT

### X-ray crystallography

For crystallization, *V. polyspora* Hop1^317-535^ was exchanged into crystallization buffer (20 mM Tris-HCl pH 7.5, 50 mM NaCl, 1 mM DTT) and concentrated to 20 mg/ml. Crystals were grown by mixing concentrated protein 1:1 with well solution containing 0.1 M MES pH 6.5 and 30% (v/v) PEG 400, in hanging-drop format. A single-wavelength anomalous diffraction (SAD) dataset was collected at ALS beamline 12.3.1 (**Table S2**). Data were indexed and integrated by XDS ^61^, then scaled and converted to structure factors by AIMLESS and TRUNCATE ^61,62^. For structure determination, anomalous sites were identified by hkl2map/SHELX ^63^ and input into the Phenix Autosol pipeline 64 for phasing and automatic model building. An initial model was manually rebuilt in COOT ^65^, and refined in phenix.refine ^66^. Figures were generated with PyMOL (version 2.0; Schrödinger, LLC) or ChimeraX ^67^.

### Cryo-electron Microscopy Sample Preparation

For cryo-EM sample preparation, reconstituted nucleosomes were incubated with a 20-fold molar excess of *S. cerevisiae* Hop1322-537 in crosslinking buffer (20 mM HEPES 7.5, 50 mM NaCl) for 30 minutes on ice, brought to room temperature for 5 minutes, then crosslinked with a final concentration of 0.02% glutaraldehyde for 30 minutes at room temperature. The crosslinking reaction was quenched using 1 volume of 1 M glycine, concentrated and passed over a Superose 6 size-exclusion column (Cytiva) (**Figure S2B**), then concentrated again to 3 μM. 3.5 μL of sample was applied to a Quantifoil Cu 1.2/1.3 300 grid (Electron Microscopy Sciences), then blotted and frozen after a 1 minute incubation using a Vitrobot Mark IV (Thermo Fisher) with blot force 20, blot time 5.5 seconds, 4°C, and 100% humidity.

### Cryo-EM Data Collection and Refinement

A full dataset of 1,314 micrographs were collected using a FEI Titan Krios G3 at magnification 130,000x (1.1 Å per pixel), using a K2 Summit direct electron detector, at a defocus range of −0.5 to −2 μm. cryoSPARC (Structura Biotechnology) version 3 was used for patch motion correction, patch CTF estimation, and particle picking (blob picker; 1,267,442 initial particles; **Figure S3**). Multiple rounds of 2D class averaging resulted in a refined set of 373,963 particles, which were re-extracted and subjected to ab initio 3D reconstruction with three classes. One class (94,297 particles) showed one Hop1 CBR bound to a nucleosome, and a second class (114,228 particles) showed a nucleosome without Hop1 bound. Both sets of particles were subjected to heterogeneous refinement, yielding final maps at 2.74 Å resolution (Hop1 CBR:nucleosome complex) and 2.57 Å (nucleosome alone) (**Table S3**). The Hop1 CBR:nucleosome complex was also subjected to local refinement using a mask covering Hop1 CBR, to a resolution of 3.15 Å (**Figure S3**). This map was used to manually rebuild a threaded model of *S. cerevisiae* Hop1^322-524^ generated by the PHYRE2 server ^68^ based on the structure of *V. polyspora* Hop1^319-529^. For the nucleosome core particle, PDB ID 3LZ0 ^69^ was used as a template for rebuilding. Models were manually rebuilt in COOT ^65^ using cryo-EM maps sharpened using DeepEMhancer ^70^, then subjected to real-space refinement in phenix.refine ^66^. To choose the correct DNA sequence orientation, both orientations of the Widom 601 sequence were built and refined, then the Pearson’s correlation coefficient between model and map were calculated for each base pair, then averaged over each orientation, and the higher-correlation orientation was used.

### Biochemical assays

For initial binding assays with *S. cerevisiae* Hop1^322-537^ and nucleosomes, biotinylated *H. sapiens* nucleosomes were purchased from Epicypher (product # 16-0006). Ten μg of *S. cerevisiae* Hop1^322-537^ was mixed with 10 μg nucleosomes in binding buffer (20 mM Tris-HCl pH 7.5, 2 mM β-mercaptoethanol, 25 mM NaCl, 10% glycerol) in 50 μL total reaction volume, and incubated at room temperature for 30 minutes. 10 μL of streptavidin beads (Dynabeads MyOne Streptavidin T1; Thermo Fisher) were added and the mixture incubated a further 30 minutes at room temperature with agitation. Beads were washed 3x with 1 mL of binding buffer, and bound proteins were eluted with SDS-PAGE loading dye, boiled, and analyzed by SDS-PAGE.

For electrophoretic mobility shift assays (EMSA), 50 μL reactions were prepared in binding buffer with nucleosome (or DNA) concentration held constant at 50 nM and varying the concentration of *S. cerevisiae* Hop1^322-537^. After 15 minute room temperature incubation, sucrose was added to a final concentration of 5% (w/v), and reactions were separated on 6% TBE-acrylamide gels pre-equilibrated (pre-run for 60 minutes at 150 V) in 0.2X TBE running buffer. Gels were run for 90 minutes at 120 Volts at 4°C. Gels were stained with Sytox Green and visualized using a Bio-Rad ChemiDoc system. Gel bands were quantified in ImageJ ^71^, and binding curves were calculated using GraphPad Prism version 9 (GraphPad Software).

>40-base DNA from Widom 601 sequence: GAGGCCGCTCAATTGGTCGTAGACAGCTCTAGCACCG

### Yeast genetics, growth, and meiotic time-course assays

*S. cerevisiae* strains were generated with PCR-based methods ^72^ or CRISPR-based mutagenesis ^73^. Strains used in this study are listed in **Table S5**. For spore viability, cells were grown on YPD agar and then patched onto SPO medium (1% KOAc) for 48–72 hours. At least 100 tetrads (400 spores) were dissected for each strain. For synchronous meiosis, cells were grown in YPD, then diluted into BYTA (1% yeast extract, 2% bactotryptone, 1% potassium acetate, 50 mM potassium phthalate) at an OD_600_ of 0.3, grown overnight, then washed and resuspended in SPO medium (0.3% potassium acetate pH 7.0) at an OD_600_ of 2.0 at 30°C to induce sporulation. Samples were taken at hour 0, 2, 3, 4, 5, 6, 7, 8 and processed. Synchrony of all time courses was verified by FACS; cells are fixed in ethanol, treated with proteinase K and RNAse prior to staining with SYTOX Green DNA staining and sorted by DNA content using BD FACSAria.

### ChIP-Sequencing

Chromatin immunoprecipitation sequencing (ChIP-seq) samples were prepared from 25mL of meiotic cultures, 3 hours after meiotic induction. Each sample was immediately crosslinked with 1% formaldehyde for 30 minutes at room temperature and then the formaldehyde was quenched by adding glycine to a final concentration of 125mM. Fixed cells were collected by centrifugation at 2000 RPM for 3 minutes, the supernatant was removed, and cell pellets were immediately frozen at −80°C. until the ChIP-seq protocol is continued. Cell pellets were resuspended in 500uL of lysis buffer (50mM HEPES/KOH pH 7.5, 140 mM NaCl, 1mM EDTA, 1% Triton X-100, 0.1% sodium deoxycholate) with protease inhibitors (1mM PMSF, 1mM Benzamidine, 1mg/ml Bacitracin, one Roche Tablet (catalog # 11836170001) in 10ml) and glass beads in a biopulveriser. Samples were sonicated at 15% amplitude for 15 seconds 5X to obtain DNA at an average length of 500bp. Sonicated cell lysate was centrifuged at 14,000 RPM for 10 minutes at 4°C to remove cell debris and the supernatant containing soluble chromatin fraction was isolated. 50uL of the chromatin fraction was set aside from each sample to use as input samples. To the remaining chromatin fraction, we added 2uL of anti-Hop1 antibody (a kind gift from Nancy Hollingsworth) and incubated rotating overnight at 4°C. Antibodies and associated DNA were then isolated using Gammabind G Sepharose beads (GE Healthcare Bio, catalog # 17-0885-01). Crosslinks were reversed by incubating samples and input at 65°C for at least 6 hours. Proteins and RNAs were removed using proteinase K and RNaseA. Libraries for ChIP sequencing were prepared by PCR amplification using Illumina TruSeq DNA sample preparation kits v1 and v2. Libraries were quality checked on 2200 Tapestation. Libraries were quantified using Qubit analysis prior to pooling. The ChIP libraries were sequenced on Illumina NextSeq 500 instruments at the NYU Biology Genomics core to yield 150 bp paired end reads. For spike-in normalization (SNP-ChIP), SK288c crosslinked meiotic samples were added to respective samples at 20% prior to ChIP processing ^42^. SNP-ChIP libraries were sequenced on NextSeq 500 to yield 150bp paired end reads. All ChIP-seq data sets are listed in **Table S6**.

### Processing of reads from Illumina sequencing

Illumina output reads were processed in the following manner. The reads were mapped to the SK1 genome (GCA_002057885.1) using Bowtie ^74^. SNP-ChIP library reads were aligned to concatenated genome assemblies of SK1 and S288c genomes ^42^. Only reads that matched perfectly to the reference genome were retrieved for further analysis. 3’ ends of the reads were extended to a final length of 200 bp using MACS2 2.1.1 (https://github.com/taoliu/MACS) and probabilistically determined PCR duplicates were removed. The input and ChIP pileups were SPMR-normalized (single per million reads) and fold-enrichment of ChIP over input data was used for further analysis. The pipeline to process Illumina reads can be found at https://github.com/hochwagenlab/ChIPseq_functions/master/ChIPseq_Pipeline_v4/. The pipeline used to process SNP-ChIP reads and calculate spike in normalization factor can be found at https://github.com/hochwagenlab/ChIPseq_functions/tree/master/ChIPseq_pipeline_hybrid_genome/.

ChIP seq datasets were made up of at least two biological replicates that were merged prior to using the ChIPseq_Pipeline_v4. All datasets were normalized to the global mean of one and regional enrichment was calculated. The R functions used can be found at https://github.com/hochwagenlab/hwglabr2/ and analysis and graphs can be found at https://github.com/hochwagenlab/Hop1-loop2.

Processed and raw data files can be found in the GEO repository: GSE225129

### Chromosome spreads

Meiotic cells were collected at hour 2, 3, and 4 after meiotic induction and treated with 200 mM Tris pH7.5/20 mM DTT for 2 min at room temperature and then spheroplasted in 2% potassium acetate/ 1 M Sorbitol/ 0.13 μg/μL zymolyase T100 at 30°C. The spheroplasts were rinsed and resuspended in ice-cold 0.1 M MES pH6.4/ 1 mM EDTA/ 0.5 mM MgCl2/ 1 M Sorbitol. Cells were placed on glass slide and one volume of fixative (1% paraformaldehyde/ 3.4% sucrose / 0.14% TritonX-100) was added to the cells on a clean glass slide (soaked in ethanol and air-dried) and distributed across the slide by gentle tilting. Four volumes of 1% lipsol were added to the slide and mixed by tilting. Cells were spread by a clean glass rod. Four volumes of fixative solution were added and slides were left to dry overnight and stored at −80°C the next day. Prior to staining, slides were washed in 1X PBS / 0.4% Kodak Photoflo, and blocked with 1X PBS / 1% chicken egg white albumin (Sigma Aldrich #A5503) on a shaker for 15 minutes at room temperature. Slides were hybridized with antibodies at 37°C for one hour (antibodies and corresponding dilutions are indicated in **Table S8**). Slides were then washed with 1X PBS/ 0.05% Triton-X three times for 5 minutes each. Secondary antibodies were hybridized at 37°C for one hour (**Table S8**). Slides were washed once with 1X PBS/ 0.05% Triton-X, once with 1X PBS/ 0.4% Kodak Photoflo, and twice with dH20 / 0.4% Kodak Photoflo and air dried prior to mounting with VECTASIELD with DAPI mounting medium (VWR #H-1200-10) and a VWR Micro Cover Glass, No. 1; 60 × 24 mm (VWR #48393 106). Stained slides were stored at 4°C prior to imaging.

### Microscopy and Cytological Analysis

Images were collected on a Deltavision Elite imaging system (GE) equipped with an Olympus 100X/1.40 NA UPLSAPO PSF oil immersion lens and an InsightSSI Solid State Illumination module. Images were captured using an Evolve 512 EMCCD camera in the conventional mode and analyzed using ImageJ software. ImageJ analysis scripts can be found at https://github.com/hochwagenlab/Hop1-loop2. Scatterplots were generated using the GGplot2 package in R.

### Western Blot Analysis

For protein extraction, 5mL of synchronous meiotic culture from time points indicated. Cells were collected and resuspended in 5mL of 5% trichloroacetic acid and incubated on ice for 10 minutes. Cells were collected by centrifugation, supernatant was poured off, and protein was transferred to an eppendorf tube with 500uL of 1M non-pHed Tris. Protein was collected by centrifugation, supernatant was poured off, and protein was resuspended in Tris-EDTA/250mM DTT. SDS loading dye was added prior to boiling samples for 10 minutes. 5uL of protein sample was loaded onto a 4-15% Mini-PROTEAN TGX Precast polyacrylamide gel from Bio-Rad (4561086). Protein samples were resolved and transferred to a PVDF membrane using the Trans-Blot STurbo Transfer System. Membranes were blocked with 5% milk /TBST solution and then hybridized to antibodies indicated (**Table S8**). Anti-Rabbit-HRP from Kindle Biosciences (R1005) from Kindle Biosciences were used with 1-shot Digital-ECL substrate (R1003) to visualize protein. Images were taken with a digital camera.

### 1D Recombination analysis

Samples were collected at the time points indicated and cells killed with 0.1% sodium azide. For pulsed field gel electrophoresis, cells wre pelleted and embedded into low melting point agarose plugs. Genomic DNA was prepared within the plugs using zymolyase and proteinase K (Roche) and resolved in a 1% LE agarose gel in 0.5x TBE using a CHEF-DRII instrument (BioRad) using the following settings: ChrVIII: 5s - 45s ramp, 5.4V/cm, 32h, 14°C; whole genome: 60s switch time for 15h followed by 90s switch time for 9h, 6V/cm, 14°C. For the analysis of specific hotspots, DNA was extracted from 10ml meiotic culture and digested with XhoI (*HIS4-LEU2*) or HindIII (*GAT1*). Fragments were separated in a 0.8% agarose gel (Seakem LE agarose/1xTBE). Gels were then washed in ethidium bromide solution to visualize DNA fragments and imaged on a Biorad Gel Doc.

For Southern blot analysis, gels were transferred to a Hybond-XL nylon membrane (GE Healthcare) using alkaline capillary transfer. Probes (available in **Table S7**) were labeled with ^32^P-dCTP using the Prime-It RT random labeling kit (Agilent, catalog #300329) and hybridized to the membrane. The membrane was then exposed using a Fuji imaging screen and phospho-signal was detected on Typhoon FLA 9000 (GE). Signal quantifications were performed using ImageJ software. All experiments were performed at least twice. Graphs were constructed using Microsoft Excel.

### 2D Recombination analysis

10ml samples were collected at time points indicated and crosslinked with psoralen (0.1mg/ml) and UV light to preserve secondary structures. Following extraction of genomic DNA, samples were digested with XhoI. Samples were separated in the first dimension in a 0.4% gel in 1xTBE and run in 2.5 L 1 ×TBE at 40V for 18 hours. The gel was equilibrated in 1xTBE containing 0.5 μg/ml ethidium bromide. Lanes were cut out, positioned perpendicular, embedded in 0.8% agarose and run in 1 ×TBE containing 0.5 μg/ml ethidium bromide at 90V for 15h. DNAs was then transferred to a nylon membrane for Southern blot analysis as described for the 1D recombination analyses.

### Evolutionary analysis of meiotic HORMAD proteins

To generate HMM profiles for the Hop1 HORMA domain and CBR (PHD, wHTH, and HTH-C domains), we used hmmbuild from the hmmer package (version HMMER 3.1b1) ^75^ based on multiple sequence alignments (MAFFT, v.7.149b ^76^ “einsi” or “linsi”) of *Saccharomycetes*-specific homologs. Two strategies were used to identify highly divergent domains: (1) Saccharomycetaceae domain-specific HMM profiles were used to query a dataset of new and previously identified HORMADs containing 147 proteins identified among 158 diverse eukaryotic genomes ^51,52^ (**Table S4**). (2) For multiple different seed sequences with the N-terminal HORMA domain removed, we ran iterative searches using jackhmmer without any heuristic filters (option ‘--max’). Resulting alignments were manually inspected for the potential presence of divergent domains (especially for wHTH). Domain identification from HMM profiles was confirmed by visual inspection of AlphaFold 2 and/or Omegafold structure predictions of each identified protein ^54^. All AlphaFold 2 structure predictions shown in **Figure 5B** were downloaded from the AlphaFold Protein Structure Database (https://www.alphafold.ebi.ac.uk) ^53^ version November 1, 2022 (AF-[Uniprot ID]-F1_model_v4.pdb files) or generated with AlphaFold Monomer version 2.0 or Omegafold using ColabFold ^77^.

## Acknowledgements

We thank N. Hollingsworth and A. MacQueen for generously sharing antibodies. CM acknowledges support from the NIH (F32 GM139386). SU acknowledges support from the UCSD Molecular Biophysics Training Grant (NIH T32 GM008326) and a National Science Foundation Graduate Research Fellowship program. ET acknowledges support from the Dutch Science Organisation (VI.Veni.202.223). AH acknowledges support from the NIH (R01 GM111715 and R35 GM148223). KDC acknowledges support from the NIH (R35 GM144121). We thank Olivia Micci-Smith, Nicole Adamski, Carolina Thornton, and everyone else at the NYU Center for Genomics and Systems Biology Genomics Core for sequencing experiments, as well as help troubleshooting the FACS experiments. This work was supported in part through the NYU IT High Performance Computing resources, services and staff expertise. The authors acknowledge the facilities of the CryoEM Facility at UC San Diego, and technical assistance of R. Ashley on cryoEM sample preparation and data collection. This work was partially conducted at the Advanced Light Source (ALS), a national user facility operated by Lawrence Berkeley National Laboratory on behalf of the Department of Energy, Office of Basic Energy Sciences, through the Integrated Diffraction Analysis Technologies (IDAT) program, supported by DOE Office of Biological and Environmental Research. Additional support comes from the National Institute of Health project ALS-ENABLE (P30 GM124169) and a High-End Instrumentation Grant S10OD018483.

## Author Contributions

CM performed ChIP-Seq experiments and data analysis, recombination analysis, and immunofluorescence, and contributed to manuscript writing and figure preparation. SU performed X-ray crystallography and cryoEM experiments, and in vitro DNA binding experiments. YG performed cryoEM sample preparation and structure determination. ET performed the evolutionary bioinformatics analyses and contributed to figure preparation and manuscript writing for this part. JZ performed immunofluorescence experiments and analysis. AH performed recombination analysis, contributed to manuscript writing and figure preparation, and obtained funding. KDC performed structure analysis, bioinformatics analysis, contributed to manuscript writing and figure preparation, and obtained funding.

