## Supplementary material for "Chromatin binding by HORMAD proteins regulates meiotic recombination initiation": Supp_Fig_1-6

### Supplementary Figures

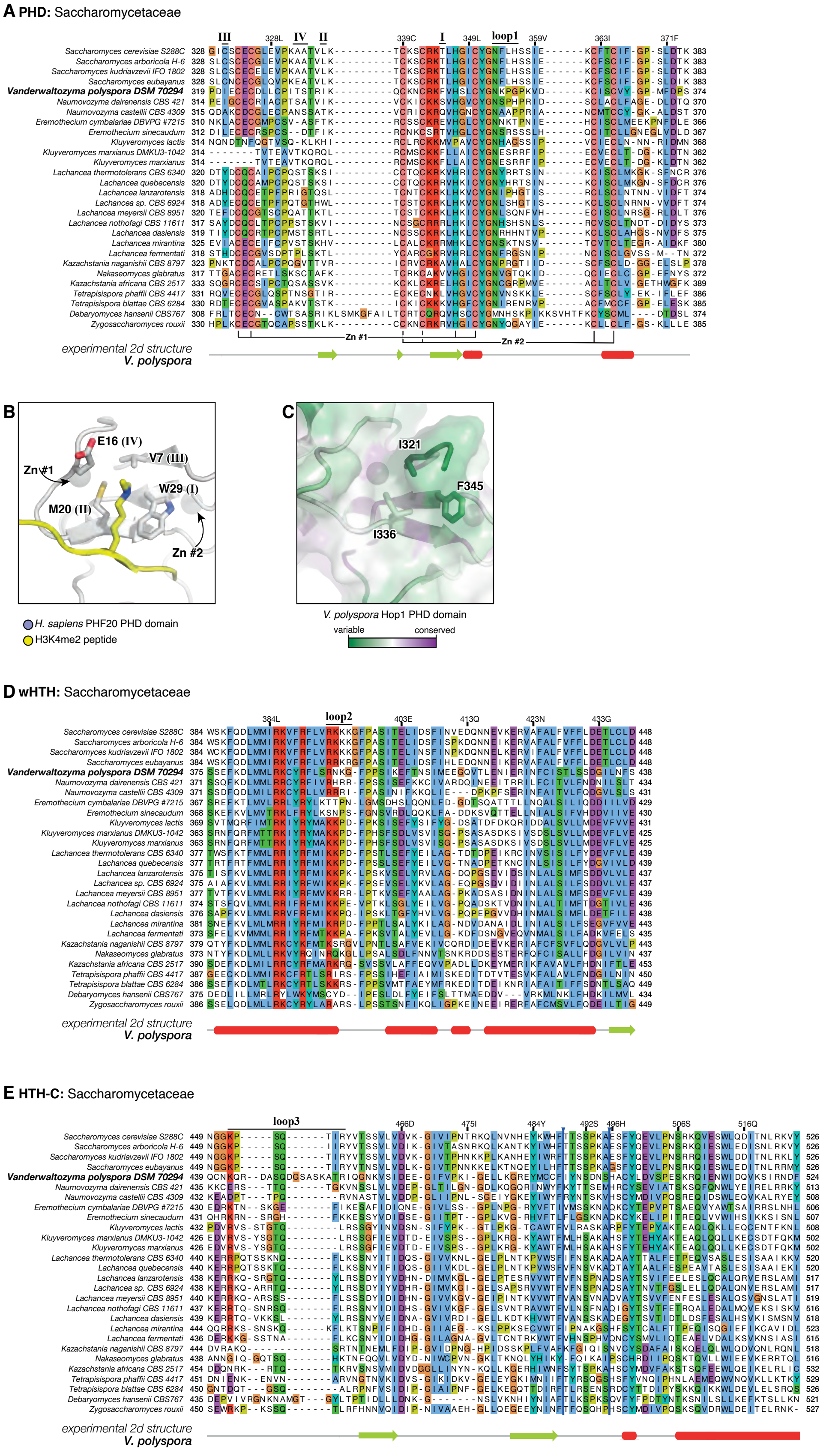
A PHD: Saccharomycetaceae
III
IV
II
I
loop1
Zn #1
Zn #2
experimental 2d structure
V. polyspora
B
E16 (IV)
V7 (III)
Zn #1
W29 (I)
M20 (II)
Zn #2
H. sapiens PHF20 PHD domain
H3K4me2 peptide
C
I321
F345
I336
V. polyspora Hop1 PHD domain
variable
conserved
D wHTH: Saccharomycetaceae
loop2
experimental 2d structure
V. polyspora
E HTH-C: Saccharomycetaceae
loop3
experimental 2d structure
V. polyspora

**Figure S1. Structure of the Hop1 CBR PHD-wHTH-HTH-C domains from Saccharomycetaceae**

(A) Sequence alignment of 28 unique Saccharomycetaceae Hop1 CBR PHD domains (NCBI accession numbers listed in **Table S2**), with residues coordinating zinc ions #1 and #2 noted at bottom, and the equivalent residues of the canonical PHD domain hydrophobic cage positions I-IV noted [39]. DNA binding loop 1 is noted at top. *V. polyspora* is used as a reference.

(B) Structure of the *H. sapiens* PHF20 PHD domain (white) bound to an H3K4me2 peptide (yellow) (PDB ID 5TBN) [79]. Shown in sticks and labeled are PHD domain positions I-IV.

(C) Structure of the *V. polyspora* Hop1 CBR PHD domain, colored by conservation within Saccharomycetaceae (green: variable; purple: conserved) as calculated by the CONSURF server [80] from the sequence alignment in panel (A). Residues corresponding to PHD domain positions I-III are shown in sticks and labeled.

(D) Sequence alignment of 28 unique Saccharomycetaceae Hop1 CBR wHTH domains (similar to panel A), with DNA binding loop 2 is noted at top. *V. polyspora* is used as a reference.

(E) Sequence alignment of 28 unique Saccharomycetaceae Hop1 CBR HTH-C domains (similar to panel A), with DNA binding loop 3 is noted at top. *V. polyspora* is used as a reference.

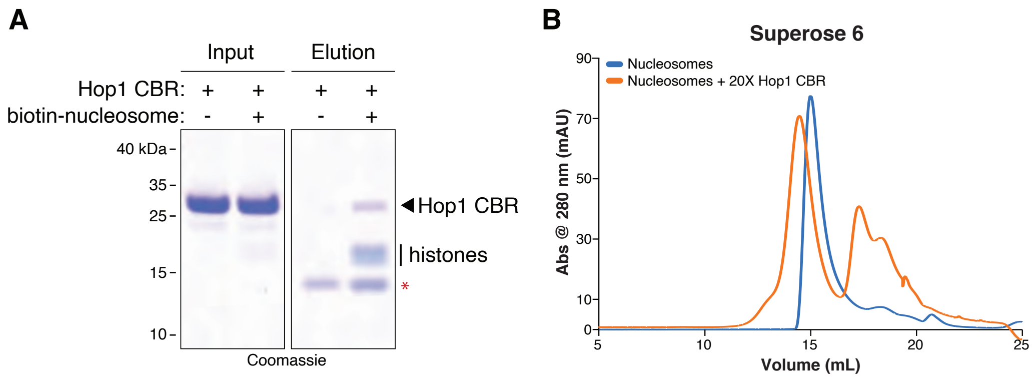
A
Input
Elution
Hop1 CBR: + + + +
biotin-nucleosome: - + - +
40 kDa
35
25
15
10
Hop1 CBR
histones
*
Coomassie
B
Superose 6
Nucleosomes
Nucleosomes + 20X Hop1 CBR
Abs @ 280 nm (mAU)
90
70
50
30
10
0
5
10
15
20
25
Volume (mL)

**Figure S2. Assembly of a Hop1 CBR:nucleosome complex**

(A) Pulldown assay with biotinylated mononucleosomes and the *S. cerevisiae* Hop1 CBR. Red asterisk indicates streptavidin eluted from the affinity resin.

(B) Superose 6 size exclusion chromatography of glutaraldehyde-crosslinked nucleosomes, either alone (blue) or pre-incubated with a 20x excess of the *S. cerevisiae* Hop1 CBR (orange).

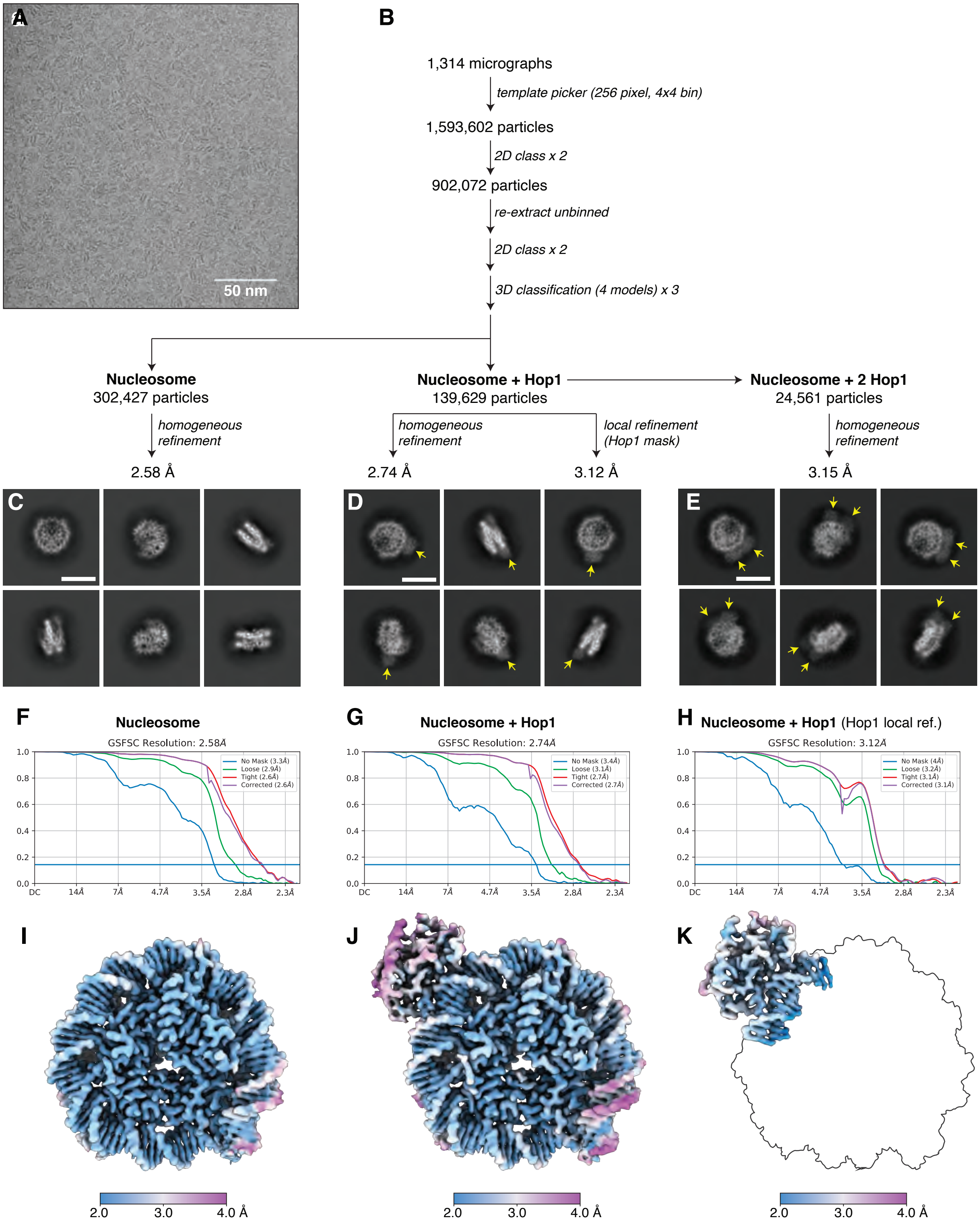

A
50 nm
B
1,314 micrographs
template picker (256 pixel, 4x4 bin)
1,593,602 particles
2D class x 2
902,072 particles
re-extract unbinned
2D class x 2
3D classification (4 models) x 3
Nucleosome
302,427 particles
homogeneous refinement
2.58 Å
Nucleosome + Hop1
139,629 particles
homogeneous refinement
2.74 Å
local refinement (Hop1 mask)
3.12 Å
Nucleosome + 2 Hop1
24,561 particles
homogeneous refinement
3.15 Å
C
D
E
F
Nucleosome
GSFSC Resolution: 2.58Å
No Mask (3.3Å)
Loose (2.9Å)
Tight (2.6Å)
Corrected (2.6Å)
G
Nucleosome + Hop1
GSFSC Resolution: 2.74Å
No Mask (3.4Å)
Loose (3.1Å)
Tight (2.7Å)
Corrected (2.7Å)
H
Nucleosome + Hop1 (Hop1 local ref.)
GSFSC Resolution: 3.12Å
No Mask (4Å)
Loose (3.2Å)
Tight (3.1Å)
Corrected (3.1Å)
1.0
0.8
0.6
0.4
0.2
0.0
DC
14Å
7Å
4.7Å
3.5Å
2.8Å
2.3Å
I
J
K
2.0
3.0
4.0 Å

**Figure S3. Cryo-EM structure of a Hop1 CBR-bound nucleosome**

(A) Raw micrograph of the Hop1 CBR-nucleosome complex. Scale bar: 50 nm.

(B) Workflow for cryo-EM structure determination of a nucleosome and the Hop1 CBR:nucleosome complex.

(C-E) Selected 2D class averages for nucleosome (C), nucleosome + Hop1 CBR (D), and nucleosome + 2 Hop1 CBR (E). In panels (D) and (E), Hop1 CBR density is indicated with yellow arrows. Scale bars = 10 nm.

(F-H) Gold-standard Fourier Shell Correlation plots for refinements of nucleosome (F), nucleosome + Hop1 CBR (G), nucleosome + Hop1 CBR (local refinement of the Hop1 CBR region) (H).

(I-K) Cryo-EM maps of nucleosome (I), nucleosome + Hop1 CBR (J), nucleosome + Hop1 CBR (local refinement of the Hop1 CBR region) (K), colored by local resolution.

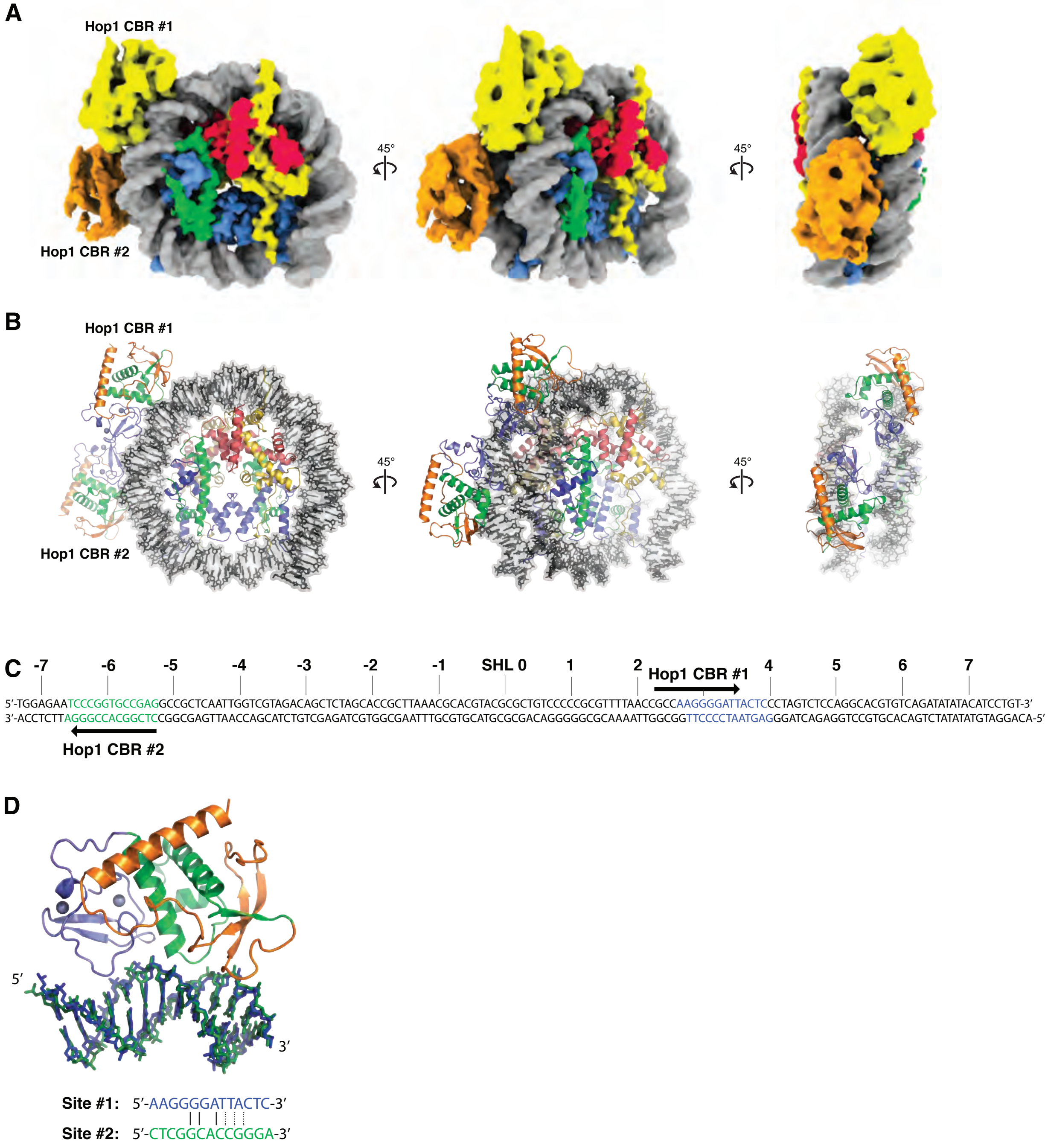
A
Hop1 CBR #1
Hop1 CBR #2
45°
45°
B
Hop1 CBR #1
Hop1 CBR #2
45°
45°
C
-7 -6 -5 -4 -3 -2 -1 SHL 0 1 2 Hop1 CBR #1 4 5 6 7
5′-TGGAGAATCCCGGTGCCGAGGCCGCTCAATTGGTCGTAGACAGCTCTAGCACCGCTTAAACGCACGTACGCGCTGTCCCCCGCGTTTTAACCGCCAAGGGGATTACTCCCTAGTCTCCAGGCACGTGTCAGATATATACATCCTGT-3′
3′-ACCTCTTAGGGCCACGGCTCCGGCGAGTTAACCAGCATCTGTCGAGATCGTGGCGAATTTGCGTGCATGCGCGACAGGGGGCGCAAAATTGGCGGTTCCCCTAATGAGGGATCAGAGGTCCGTGCACAGTCTATATATGTAGGACA-5′
Hop1 CBR #2
D
5′
3′
Site #1: 5′-AAGGGGATTACTC-3′
Site #2: 5′-CTCGGCACCGGGA-3′

**Figure S4. Structure of a nucleosome + 2 Hop1 CBR complex**

(A) Three views of cryoEM density for a nucleosome + 2 Hop1 CBR complex (3.15 Å overall resolution, gaussian smoothed with a σ of 1.1 Å). DNA is colored gray, histones colored yellow (H2A), red (H2B), blue (H3), and green (H4), and two Hop1 CBR domains colored yellow and orange.

(B) Molecular model for a nucleosome + 2 Hop1 CBR complex, colored as in panel (A) except Hop1 CBR is colored blue (PHD), green (wHTH), and orange (HTH-C).

(C) Schematic of the Widom 601 DNA sequence used for nucleosome assembly, with binding sites for Hop1 CBR #1 (blue) and #2 (green) noted. SHL: superhelical locations, 0 at the nucleosome dyad axis to 7 at the DNA ends.

(D) Structural overlay of the DNA bound to Hop1 CBR #1 (blue) and #2 (green). In sequence alignment at bottom, solid lines indicate identity and dotted lines indicate shared status as either pyrimidine or purine.

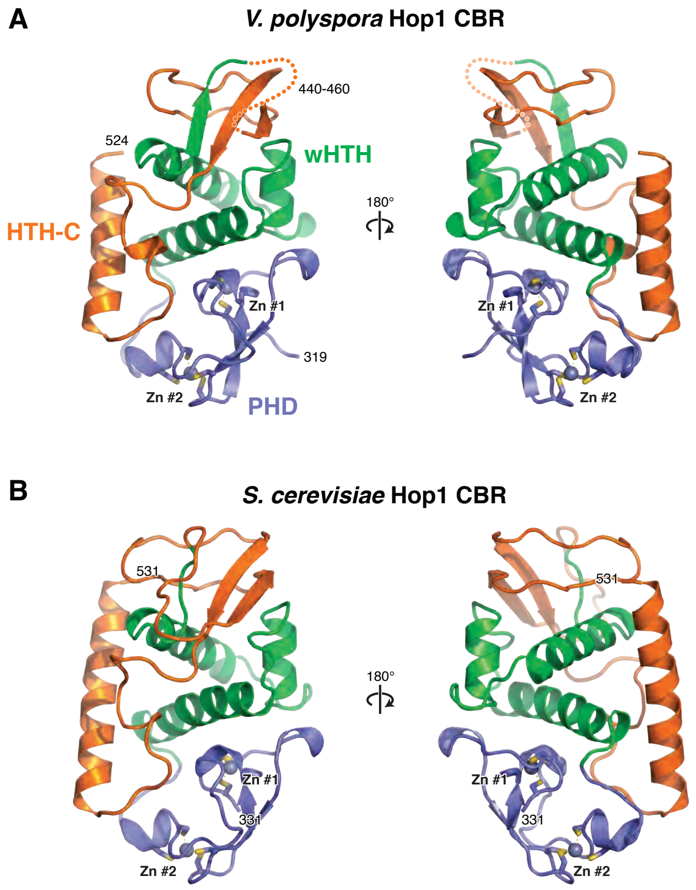

A
V. polyspora Hop1 CBR
440-460
524
wHTH
HTH-C
180°
Zn #1
319
Zn #2
PHD
B
S. cerevisiae Hop1 CBR
531
180°
Zn #1
331
Zn #2

**Figure S5. Hop1 CBR structure comparison**

(A) Two views of the *V. polyspora* Hop1 CBR crystal structure, with PHD domain colored blue, wHTH green, and HTH-C orange. Residues 440-460 in the HTH-C region are disordered and represented as a dotted line.

(B) Two views of the *S. cerevisiae* Hop1 CBR from the nucleosome-bound cryoEM structure. The two structures overlay with an overall $C_\alpha$ r.m.s.d. of 1.06 Å (111 residue pairs aligned).

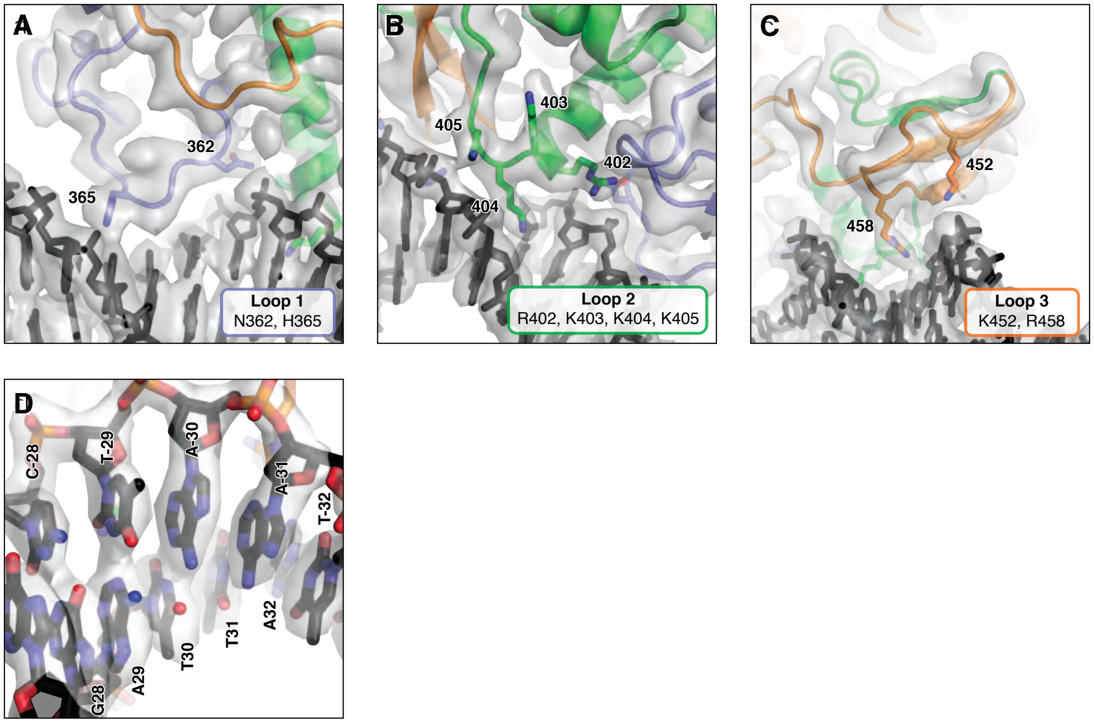
A
362
365
Loop 1
N362, H365
B
403
405
402
404
Loop 2
R402, K403, K404, K405
C
452
458
Loop 3
K452, R458
D
C-28
T-29
A-30
A-31
T-32
G28
A29
T30
T31
A32

**Figure S6. Hop1 CBR-DNA interactions**

(A-C) CryoEM density (semi-transparent surface) at 2.74 Å resolution (unsharpened) for the Hop1 CBR Loop 1 (panel A), Loop 2 (B), and Loop 3 (C) interactions with DNA. DNA-interacting residues are labeled.

(D) CryoEM density showing a portion of nucleosomal DNA within the Hop1 CBR-bound region, showing unambiguous assignment of the orientation of the Widom 601 DNA sequence. Chain I residues -28 to -32 are at top/front, and Chain J residues 28 to 32 are at bottom/back.
