## Supplementary material for "Chromatin binding by HORMAD proteins regulates meiotic recombination initiation": Supp_Fig_7-11

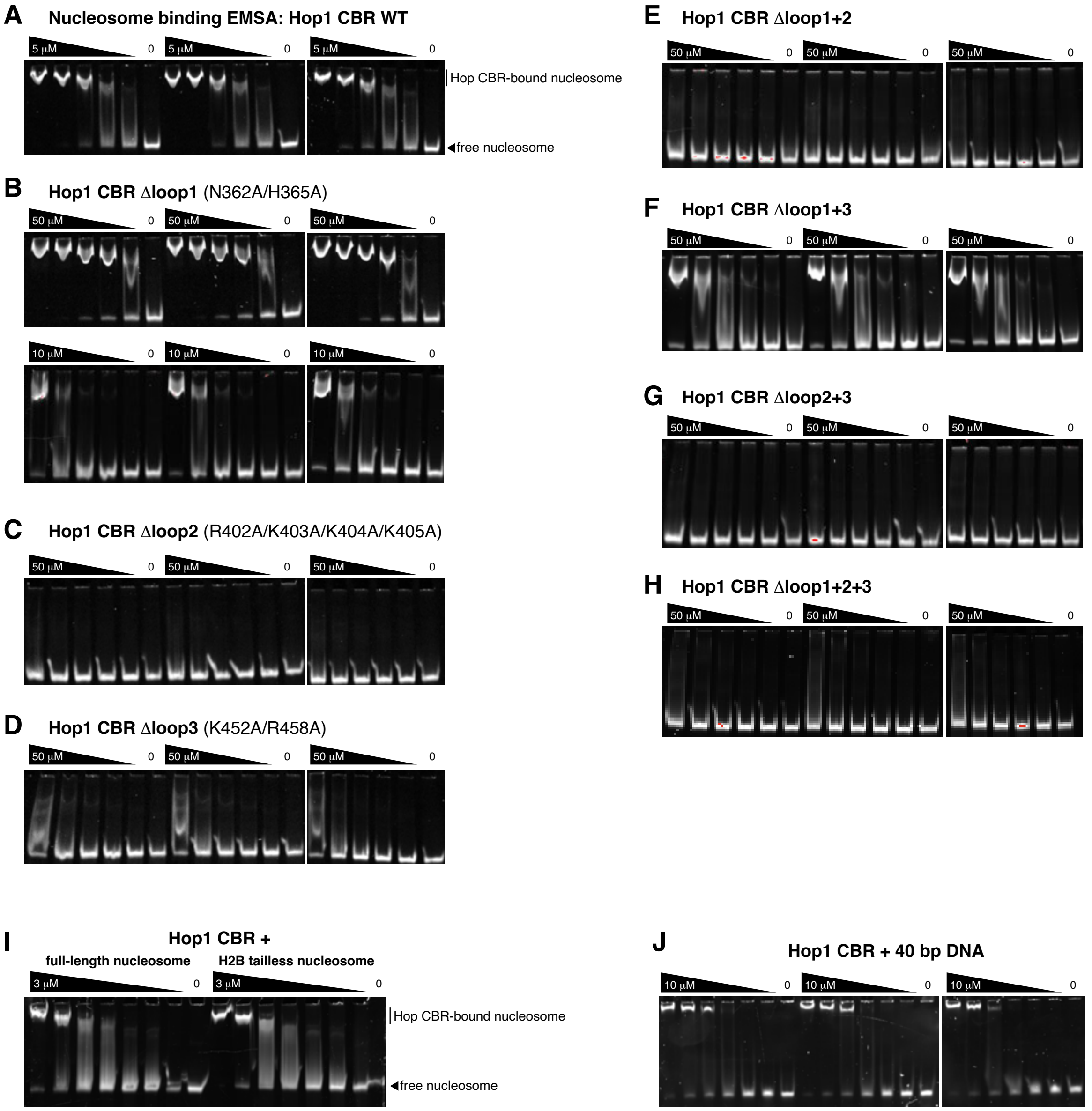
A
Nucleoside binding EMSA: Hop1 CBR WT
5 μM
0
Hop CBR-bound nucleosome
free nucleosome
B
Hop1 CBR Δloop1 (N362A/H365A)
50 μM
10 μM
0
C
Hop1 CBR Δloop2 (R402A/K403A/K404A/K405A)
50 μM
0
D
Hop1 CBR Δloop3 (K452A/R458A)
50 μM
0
E
Hop1 CBR Δloop1+2
50 μM
0
F
Hop1 CBR Δloop1+3
50 μM
0
G
Hop1 CBR Δloop2+3
50 μM
0
H
Hop1 CBR Δloop1+2+3
50 μM
0
I
Hop1 CBR +
full-length nucleosome
H2B tailless nucleosome
3 μM
0
Hop CBR-bound nucleosome
free nucleosome
J
Hop1 CBR + 40 bp DNA
10 μM
0

**Figure S7. Nucleosome and DNA binding by the *S. cerevisiae* Hop1 CBR**

(A) Triplicate electrophoretic mobility shift assays (EMSAs) for wild-type Hop1 CBR binding to reconstituted nucleosomes. The highest concentration of Hop1 CBR used was 5 µM, followed by serial 2x dilutions. Band intensities were quantified in ImageJ and $K_d$ was calculated in Prism using a single-site binding model ($K_d$ = 0.47 +/- 0.11 µM).

(B) Triplicate EMSAs for Hop1 CBR Δloop1 mutant (N362A/H365A) binding nucleosomes, with two different starting protein concentrations (top: 50 µM; bottom: 10 µM) ($K_d$ = 2.0 +/- 0.4 µM).

(C) Triplicate EMSAs for Hop1 CBR Δloop2 mutant (R402A/K403A/K404A/K405A) binding nucleosomes ($K_d$ > 50 µM).

(D) Triplicate EMSAs for Hop1 CBR Δloop3 mutant (K452A/R456A) binding nucleosomes ($K_d$ ~ 50 µM).

(E) Triplicate EMSAs for Hop1 CBR Δloop1+2 mutant binding nucleosomes ($K_d$ > 50 µM).

(F) Triplicate EMSAs for Hop1 CBR Δloop1+3 mutant binding nucleosomes ($K_d$ = 13 +/- 3 µM).

(G) Triplicate EMSAs for Hop1 CBR Δloop2+3 mutant binding nucleosomes ($K_d$ > 50 µM).

(H) Triplicate EMSAs for Hop1 CBR Δloop1+2+3 mutant binding nucleosomes ($K_d$ > 50 µM).

(I) EMSAs comparing binding of wild-type Hop1 CBR to nucleosomes reconstituted with full-length (left) or tailless (right) histone H2B.

(J) Triplicate EMSAs for wild-type Hop1 CBR binding a 40-bp DNA encompassing its preferred binding site on the Widom 601 DNA ($K_d$ = 2.1 +/- 0.6 µM).

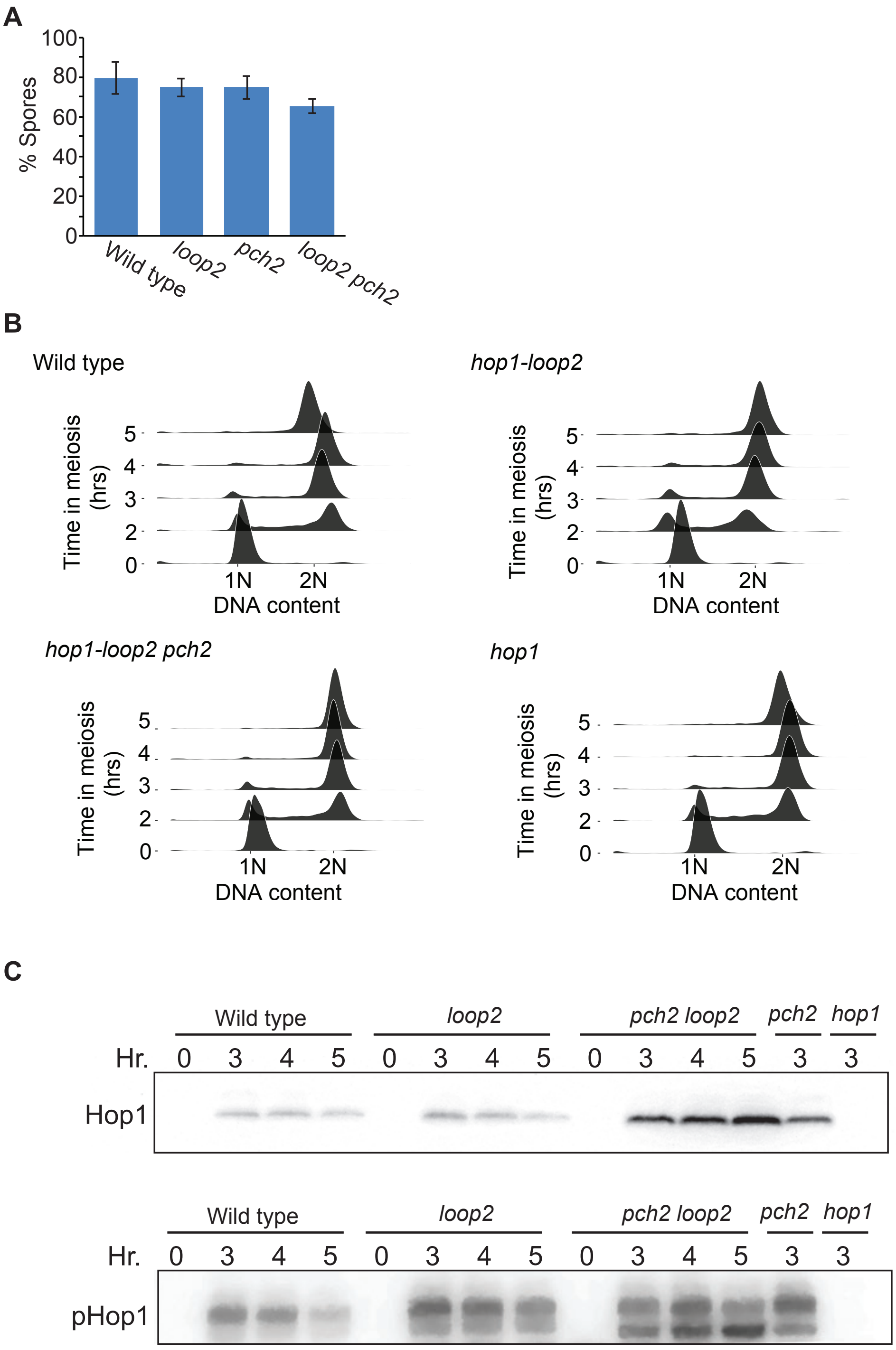
A
100
80
60
40
20
0
% Spores
Wild type
loop2
pch2
loop2 pch2
B
Wild type
hop1-loop2
hop1-loop2 pch2
hop1
Time in meiosis (hrs)
5
4
3
2
0
1N
2N
DNA content
C
Wild type
loop2
pch2 loop2
pch2
hop1
Hr.
0 3 4 5 0 3 4 5 0 3 4 5 3 3
Hop1
pHop1

**Figure S8. Hop1 protein and phosphorylation levels across genotypes**

(A) Sporulation efficiency data for the indicated genotypes.
(B) Flow cytometry analysis of DNA content, indicating synchronous meiotic entry.
(C) Western blots showing protein levels across genotypes. Top shows western blot of Hop1 protein. Bottom shows western blot using an antibody specific to phosphorylated Hop1 T318 [23].

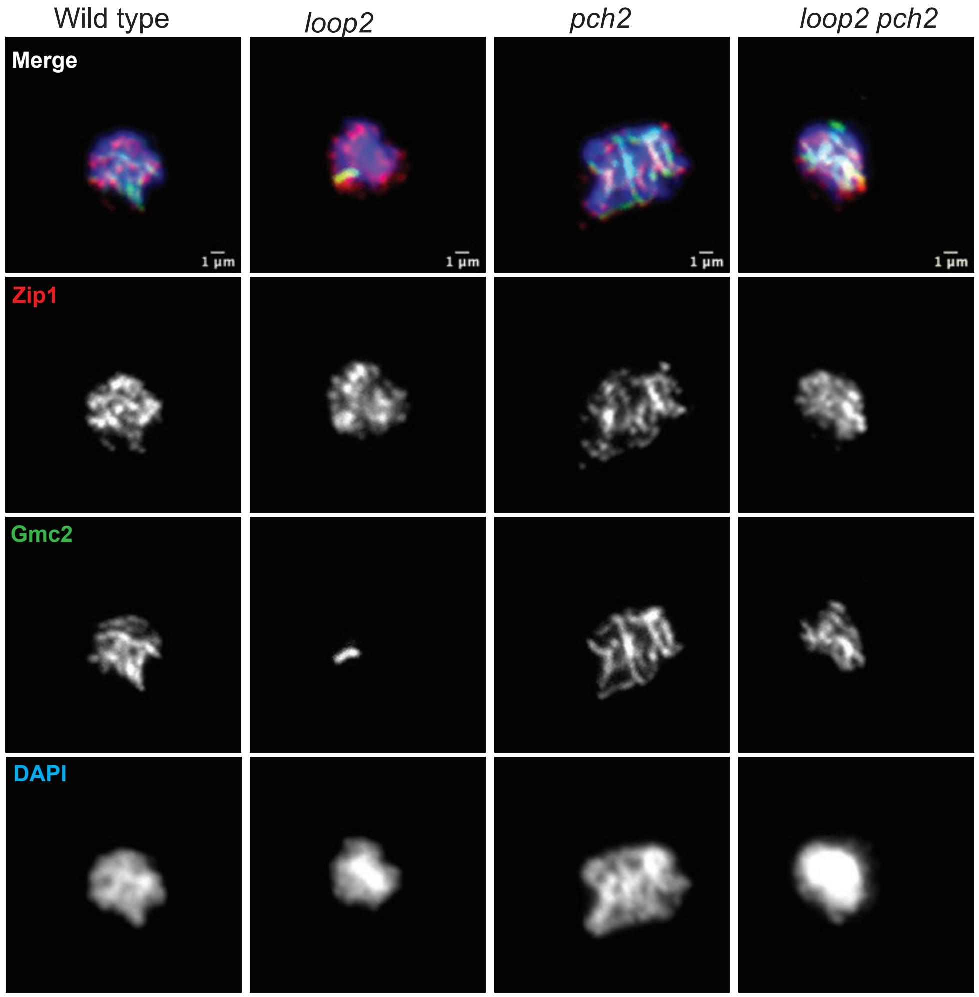
Wild type
loop2
pch2
loop2 pch2
Merge
1 μm
1 μm
1 μm
1 μm
Zip1
Gmc2
DAPI

**Figure S9. Localization of the central element protein Gmc2**

Samples were taken from meiotic cultures of the indicated genotypes at hour 3, chromosome spreads were

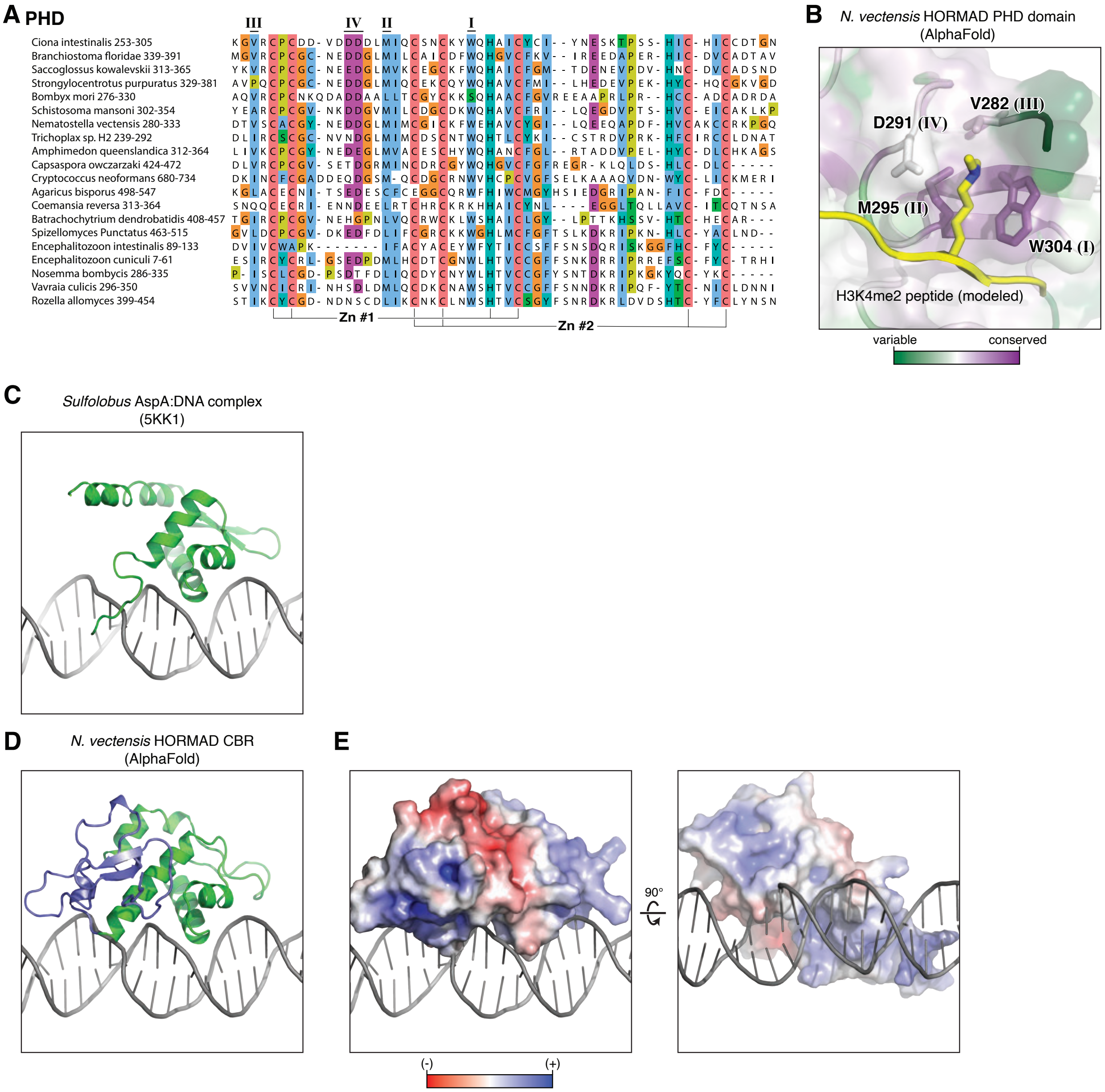
A PHD
III
IV
II
I
Ciona intestinalis 253-305
Branchiostoma floridae 339-391
Saccoglossus kowalevskii 313-365
Strongylocentrotus purpuratus 329-381
Bombyx mori 276-330
Schistosoma mansoni 302-354
Nematostella vectensis 280-333
Trichoplax sp. H2 239-292
Amphimedon queenslandica 312-364
Capsaspora owczarzaki 424-472
Cryptococcus neoformans 680-734
Agaricus bisporus 498-547
Coemansia reversa 313-364
Batrachochytrium dendrobatidis 408-457
Spizellomyces Punctatus 463-515
Encephalitozoon intestinalis 89-133
Encephalitozoon cuniculi 7-61
Nosemma bombycis 286-335
Vavraia culicis 296-350
Rozella allomyces 399-454
Zn #1
Zn #2
B
N. vectensis HORMAD PHD domain
(AlphaFold)
V282 (III)
D291 (IV)
M295 (II)
W304 (I)
H3K4me2 peptide (modeled)
variable
conserved
C
Sulfolobus AspA:DNA complex
(5KK1)
D
N. vectensis HORMAD CBR
(AlphaFold)
E
90°
(-)
(+)

**Figure S10. Meiotic HORMAD PHD domains have a conserved binding pocket**

(A) Sequence alignment of the PHD domain of meiotic HORMADs found among Opisthokonta (Fungi+animals+unicellular relatives) with a PHD+wHTH domain pair (excluding HORMADs with HTH-C) shown in **Figure 5A**, focusing on the PHD domain. Residues coordinating zinc ions #1 and #2 noted at bottom, and the equivalent residues of the canonical PHD domain hydrophobic cage positions I-IV noted [39].

(B) AlphaFold 2 model structure of *N. vectensis* meiotic HORMAD (Uniprot ID A7RLI6), focusing on the putative binding pocket for a lysine residue in a histone tail. Shown in yellow is an H3K4me2 peptide modeled from a structure of the *H. sapiens* PHF20 PHD domain (white) bound to an H3K4me2 peptide (PDB ID 5TBN) [79]. The *N. vectensis* meiotic HORMAD PHD domain is colored by conservation, as calculated by the CONSURF server [80] from the sequence alignment in panel (a).

(C) Structure of a model wHTH domain, *Sulfolobus* sp. NOBH2 pNOB8 AspA (green) bound to DNA (gray) (PDB ID 5KK1; not published)

(D) Model of the DNA-bound *N. vectensis* HORMAD CBR, created by overlaying the *N. vectensis* HORMAD CBR (PHD blue, wHTH green) onto AspA.

(E) Two views of the DNA-bound *N. vectensis* HORMAD CBR model, shown with electrostatic surface calculated by APBS [81]. The surface predicted to bind DNA is positively charged (blue).

**wHTH:** Eukarya

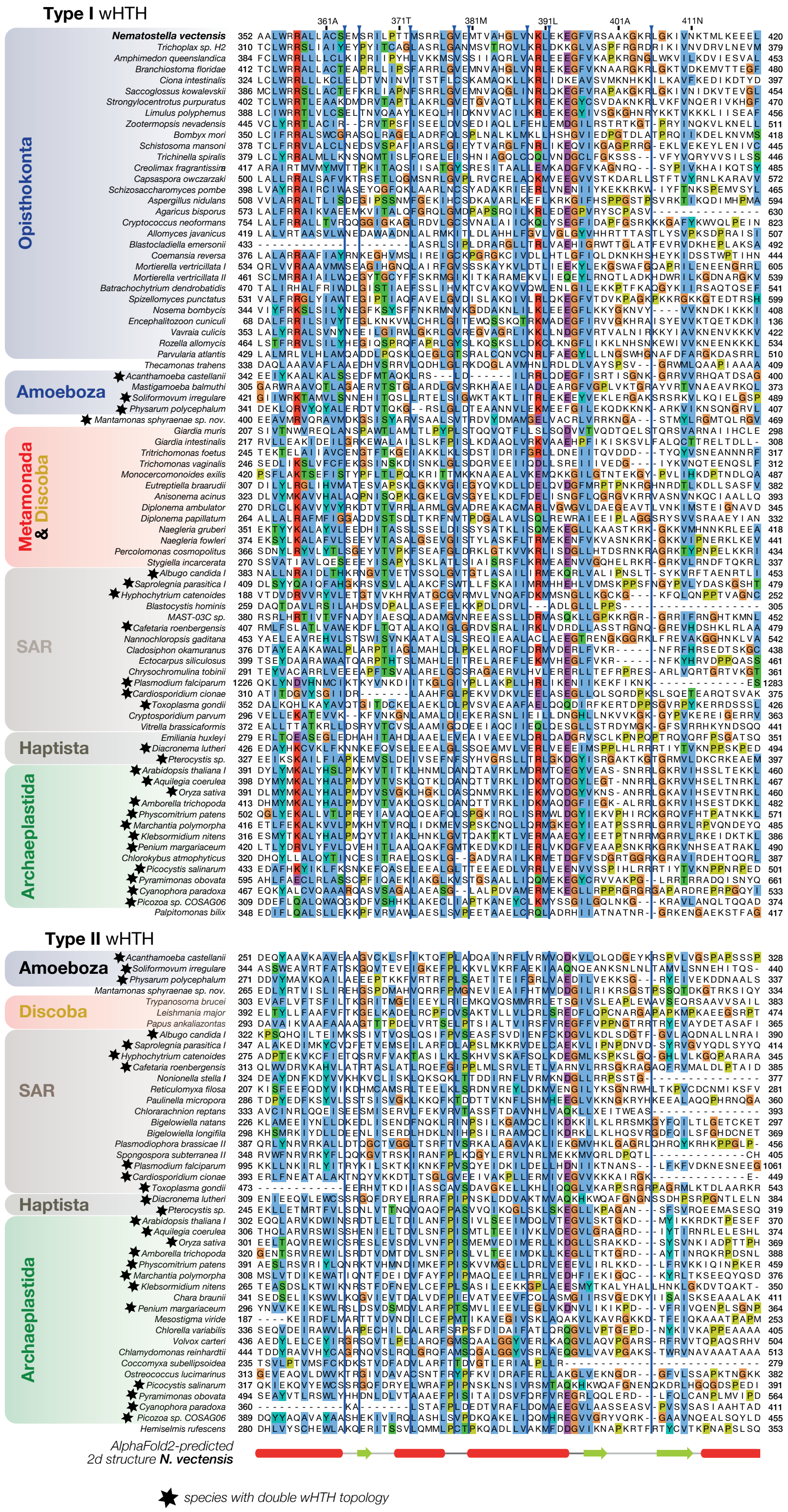

**Figure S11. Meiotic HORMAD wHTH domains can be classified into two types that are each present in meiotic HORMADs with a tandem wHTH configuration**

Sequence alignment of wHTH domains found in 75 single wHTH and 25 double wHTH among 105 different eukaryotic species (**Table S4**) - exclude those of the Saccharomycetaceae. To make the alignment more easily readable, we used the wHTH domain of *Nematostella vectensis* (Uniprot ID A7RLI6) as a reference and excluded any column in the multiple alignment that was not found in this domain (see blue lines for marks of column removal). Full multiple alignments can be found in **File S1**. wHTH domains were manually classified into two types that correspond to either one of the wHTH domains found in meiotic HORMADs with two wHTH domains (see star). Phylogenetic tree analyses did not yield statistically sensible trees, likely due to the rather high sequence divergence found among the wHTH domains in these meiotic HORMAD proteins. Type 1 wHTHs are characterized by a conserved positively charged patch in the first alpha helix, while Type 2 wHTHs harbor a conserved ‘FP’ motif in a loop between two central alpha helices (see for classification **Table S4**). Note that the wHTH domain is conserved until the first beta strand of the ‘wing’. Most wHTHs in other meiotic HORMADs found among eukaryotes do have a second strand and a capping helix, but this part of the domain is highly divergent between lineages (i.e. where loop 3 in Saccharomycetaceae resides).
