## Supplementary material for "Chromatin binding by HORMAD proteins regulates meiotic recombination initiation": Supp_Tables

### Supplementary Information

#### Supplementary Tables

##### Table S1. Crystallographic data collection and refinement

| **Data collection** | ***Vp* Hop1 Zn SAD** |
|---|---|
| Synchrotron/Beamline | ALS 12.3.1 |
| Date collected | May 14, 2015 |
| Resolution (Å) | 32.6 – 1.51 |
| Wavelength (Å) | 1.283 |
| Space Group | $P2_1$ |
| Unit Cell Dimensions (a, b, c) Å | 46.40, 38.94, 69.00 |
| Unit cell Angles (α,β,γ) ° | 90, 109.05, 90 |
| $I/\sigma$ (last shell) | 13.9 (1.2) |
| [a] $R_{sym}$ (last shell) | 0.048 (0.729) |
| [b] $R_{meas}$ (last shell) | 0.076 (0.628) |
| [c] $CC_{1/2}$, last shell | 0.53 |
| Completeness (last shell) % | 92.2 (61.0) |
| Number of reflections | 219746 |
| *unique* | 65701 |
| Multiplicity (last shell) | 3.3 (2.0) |
| Number of sites | 2 |
| **Refinement** | |
| Resolution (Å) | 32.6 – 1.55 |
| No. of reflections | 32716 |
| *working* | 30347 |
| *free* | 2369 |
| [e] $R_{work}$ (%) | 15.47 |
| [e] $R_{free}$ (%) | 18.08 |
| **Structure/Stereochemistry** | |
| Number of atoms | 3237 |
| *hydrogen* | 1515 |
| *solvent* | 210 |
| r.m.s.d. bond lengths (Å) | 0.017 |
| r.m.s.d. bond angles (°) | 1.596 |
| Ramachandran favored/allowed | 98.91%/100.0% |
| Poor rotamers | 0.57% |
| MolProbity Score | 1.28 |
| Clashscore (all atoms) | 5.25 |
| [f] PDB ID | 7UBA |
| [g] SBGrid Data Bank ID | 826 |

[a] $R_{sym} = \sum\sum_j |I_j - \langle I \rangle| / \sum I_j$, where $I_j$ is the intensity measurement for reflection j and $\langle I \rangle$ is the mean intensity for multiply recorded reflections.

[b] $R_{meas} = \sum_h [\sqrt{(n/(n-1))} \sum_j [I_{hj} - \langle I_h \rangle] / \sum_{hj} \langle I_h \rangle$

where $I_{hj}$ is a single intensity measurement for reflection h, $\langle I_h \rangle$ is the average intensity measurement for multiply recorded reflections, and $n$ is the number of observations of reflection h.

[c] $CC_{1/2}$ is the Pearson correlation coefficient between the average measured intensities of two randomly-assigned half-sets of the measurements of each unique reflection.

[e] $R_{work,\ free} = \sum ||F_{obs}| - |F_{calc}|| / |F_{obs}|$, where the working and free *R*-factors are calculated using the working and free reflection sets, respectively.

[f] Coordinates and structure factors have been deposited in the RCSB Protein Data Bank (http://www.rcsb.org).

[g] Diffraction data have been deposited with the SBGrid Data Bank (http://data.sbgrid.org) with the noted accession codes.

#### Table S2. Saccharomycetaceae Hop1 proteins used for sequence alignments

| NCBI Accession # | Species |
|---|---|
| NP_012193.3 | *Saccharomyces cerevisiae* S288C |
| EJS43276.1 | *Saccharomyces arboricola* H-6 |
| EJT42208.1 | *Saccharomyces kudriavzevii* IFO 1802 |
| XP_018221544.1 | *Saccharomyces eubayanus* |
| XP_001642921.1 | *Vanderwaltozyma polyspora* DSM 70294 |
| XP_003667640.1 | *Naumovozyma dairenensis* CBS 421 |
| XP_003674303.1 | *Naumovozyma castellii* CBS 4309 |
| XP_003648139.1 | *Eremothecium cymbalariae* DBVPG#7215 |
| XP_017987107.1 | *Eremothecium sinecaudum* |
| XP_452539.1 | *Kluyveromyces lactis* |
| BAO40435.1 | *Kluyveromyces marxianus* DMKU3-1042 |
| BAP71922.1 | *Kluyveromyces marxianus* |
| XP_002551673.1 | *Lachancea thermotolerans* CBS 6340 |
| CUS23296.1 | *Lachancea quebecensis* |
| CEP60095.1 | *Lachancea lanzarotensis* |
| SCU79936.1 | *Lachancea* sp. CBS 6924 |
| SCU95736.1 | *Lachancea meyersii* CBS 8951 |
| SCU82479.1 | *Lachancea nothofagi* CBS 11611 |
| SCU91559.1 | *Lachancea dasiensis* |
| *SCU92480.1* | *Lachancea mirantina* |
| *SCV99872.1* | *Lachancea fermentati* |
| CCK69524.1 | *Kazachstania naganishii* CBS 8797 |
| XP_449898.1 | *Candida glabrata* |
| XP_003955564.1 | *Kazachstania africana* CBS 2517 |
| XP_003688665.1 | *Tetrapisispora phaffii* CBS 4417 |
| *XP_004182647.1* | *Tetrapisispora blattae* CBS 6284 |
| XP_459121.2 | *Debaryomyces hansenii* CBS767 |
| XP_002496029.1 | *Zygosaccharomyces rouxii* |

#### Table S3. Cryo-electron microscopy data collection and refinement

| Data collection | |
|---|---|
| Microscope | TFS Titan Krios G3 |
| Voltage (keV) | 300 |
| Nominal magnification | 130,000x |
| Exposure navigation | Image Shift |
| Cumulative Exposure (e-/Å$^2$) | 50.02 |
| Exposure rate (e-/pixel/sec) | 6 |
| Detector | Gatan K2 |
| Pixel size (Å) | 1.1 |
| GIF slit width (eV) | 20 |
| Defocus range (µm) | -0.5 to -2 |
| Micrographs collected | 1,314 |

| Reconstruction | Hop1 CBR + Nucleosome | Nucleosome |
|---|---|---|
| Final particles (no.) | 139,629 | 302,427 |
| B-factor (Å$^2$) | 75.0 | 80.7 |
| Resolution (Å) | | |
| FSC 0.5 (unmasked/masked) | 4.51/3.18 | 3.99/3.01 |
| FSC 0.143 (unmasked/masked) | 3.45/2.74 | 3.29/2.78 |
| [c]Resolution range (25th/75th percentile) | 2.62-4.92 | 2.46-4.79 |

| Refinement | Hop1 CBR + Nucleosome | Nucleosome |
|---|---|---|
| Number of atoms | 13625 | 12029 |
| *ligands* | 2 (Zn) | 0 |
| Model-Map Correlation Coefficient (masked) | 0.87 | 0.86 |
| Model-Map Resolution (Å) | | |
| FSC 0.5 (unmasked/masked) | 3.2/3.1 | 3.0/2.9 |
| FSC 0.143 (unmasked/masked) | 2.8/2.7 | 2.6/2.6 |
| r.m.s.d. bond lengths (Å) | 0.003 | 0.003 |
| r.m.s.d. bond angles (°) | 0.479 | 0.552 |
| Ramachandran (%) | | |
| Outliers | 0 | 0 |
| Allowed | 2.65 | 1.87 |
| Favored | 97.35 | 98.13 |
| Poor rotamers (%) | 0.25 | 0.32 |
| MolProbity Score | 0.88 | 0.89 |
| Clashscore (all atoms) | 0.83 | 1.52 |
| EMRinger Score | 3.79 | 3.06 |
| [a] PDB ID | 8CWW | 8CZE |
| [b] EMDB ID | 27030 | 27096 |

[a] Coordinates have been deposited in the RCSB Protein Data Bank (http://www.rcsb.org).

[b] EM density maps (final unsharpened and sharpened maps, half maps, and masks) have been deposited to the Electron Microscopy Data Bank (https://pdbe.org/emdb).

[C] Local resolution range calculated at atom positions from final model.

**Table S4. Meiotic HORMAD protein domains and sequences from 158 diverse eukaryotes**

*(see attached Excel workbook)*

**Table S5. Yeast strains used in this study**

| strain | genotype | reference |
|---|---|---|
| H7797 | MATa/MATalpha, ho::LYS2/ho::LYS2, lys2/lys2, ura3/URA3, leu2::hisG/LEU2, his3::hisG/HIS3, trp1::hisG/TRP1 | PMID: 21376234 |
| H8644 ("SK288c") | MATa/MAT alpha, his3Δ1/HIS3, LEU/leu2Δ0, LYS/lys2Δ0, ura3Δ0/URA3, RME1(ins-308a)/RME1(ins-308a), TAO3(E1493Q)/TAO3(E1493Q), MKT1(D30G)/MKT1(D30G) | PMID: 21816273 |
| H9120 | MATalpha/MATa, ho::LYS2/ho::LYS2, lys2/lys2, leu2::hisG/leu2::hisG, HIS/his3::hisG, ura3::hisG/URA3, trp1::hisG/TRP1, hop1::LEU2/hop1::LEU2 | |
| H11644 | MATa/MATalpha, ho::LYS2/ho::LYS2, lys2/lys2, ura3/URA3, leu2::hisG/LEU2, his3::hisG/HIS3, trp1::hisG/TRP1, hop1-loop2/hop1-loop2<br>(hop1-loop2 = R402A, K403A, K404A, K405A) | |
| H11757 | MATa/MATalpha, ho::LYS2/ho::LYS2, lys2/lys2, URA3/ura3, LEU2/leu2::hisG, his3::hisG/HIS3, trp1::hisG/TRP1, pch2Δ::KanMX, hop1-loop2/pch2Δ::KanMX, hop1-loop2 | |
| H11758 | MATalpha/MATa, ho::LYS2/ho::LYS2, lys2/lys2, URA3/ura3, LEU2/leu2::hisG, HIS3/his3::hisG, TRP1/trp1::hisG, pch2Δ::KanMX/pch2Δ::KanMX | |
| H11276 | MATa/MATalpha, ho::hisG/ho::hisG, leu2::hisG/leu2::hisG, ura3(Δsma-pst::hisG)/ura3(Δsma-pst::hisG), HIS4::LEU2-(NBam;ori)/his4X::LEU2-(NgoMIV)-URA3 | |
| H11688 | MATa/MATalpha, ho::hisG(?)/ho::hisG(?), LYS2/LYS2, leu2::hisG/leu2::hisG, ura3(Δsma-pst::hisG)/ura3(Δsma-pst::hisG), HIS3/his3::hisG(?), TRP1/TRP, his4X::LEU2-(NgoMIV)-URA3, hop1-loop2<br>HIS4::LEU2-(NBam;ori), hop1-loop2 | |
| H11811 | MATa, ho::hisG(?), LYS2, leu2::hisG, ura3(Δsma-pst::hisG), HIS3, TRP1,<br>MATalpha, ho::hisG(?), LYS2, leu2::hisG, his3::hisG(?), trp1::hisG, ura3, his4X::LEU2-(NgoMIV)-URA3, hop1-loop2/hop1-loop2, pch2Δ::KanMX/HIS4::LEU2-(NBam;ori)/pch2Δ::KanMX4 | |
| H11812 | MATa/MATalpha, ho::hisG(?)/ho::hisG(?), LYS2/LYS2, leu2::hisG/leu2::hisG, ura3(Δsma-pst::hisG)/ura3, HIS3/his3::hisG(?), trp1::hisG/TRP1, his4X::LEU2-(NgoMIV)-URA3, pch2Δ::KanMX/HIS4::LEU2-(NBam;ori), pch2Δ::KanMX4 | |
| H11569 | MATa/MATalpha, ho::hisG/ho::hisG, LYS2/LYS2, leu2::hisG/leu2::hisG, ura3(Δsma-pst::hisG)/ura3(Δsma-pst::hisG), HIS3/his3::hisG, trp1::hisG/TRP1, HIS4::LEU2-(NBam;ori)/HIS4::LEU2-(NBam;ori), rad50S::URA3/rad50S::URA3, hop1::KanMX/hop1::KanMX (KanMX replacing aa 71-134) | |
| H11570 | MATa/MATalpha, ho::hisG/ho::hisG, LYS2/LYS2, leu2::hisG/leu2::hisG, ura3(Δsma-pst::hisG)/ura3(Δsma-pst::hisG), his3::hisG/HIS3, trp1::hisG/TRP1 HIS4::LEU2-(NBam;ori)/HIS4::LEU2-(NBam;ori), rad50S::URA3/rad50S::URA3 | |
| H11810 | MATa/MATalpha, ho::LYS2/ho::LYS2, lys2/lys2, ura3/ura3, LEU2/LEU2, HIS3/HIS3, TRP1/trp1::hisG, rad50S::URA3/rad50S::URA3, pch2Δ::KanMX/pch2Δ::KanMX | |
| H11809 | MATa/MATalpha, ho::LYS2/ho::LYS2, lys2/lys2, ura3/ura3, leu2::hisG/LEU2, HIS3/HIS3, TRP1/TRP1, hop1-loop2/hop1-loop2, rad50S::URA3/rad50S::URA3, pch2Δ::KanMX/pch2Δ::KanMX | |
| H12322 | MATa/MATalpha, ho::LYS2/ho::LYS2, lys2/lys2, URA3/ura3, leu2::hisG/LEU2, HIS3/his3::hisG(?), TRP1/trp1::hisG, tel1Δ::HIS3/tel1Δ::HIS3 | |
| H12321 | MATa/MATalpha, ho::LYS2/ho::LYS2, lys2/lys2, URA3/ ura3, leu2::hisG/LEU2, his3::hisG/ his3::hisG(?), trp1::hisG/TRP1, tel1Δ::HIS3/tel1Δ::HIS3, hop1-loop2/hop1-loop2 | |

**Table S6. ChIP-sequencing Data Sets**

| Type | Genotype | File Name |
|---|---|---|
| aligned/Bedgraph | loop2 | 11644-NHantiHop1-20220331-20220125-20221004-Reps-SK1Yue-PM-PE_B4_W3_MACS2_FE.bdg.gz |
| aligned/Bedgraph | wild type | Hop1-wildtype-334-340-32-177-Reps-SK1Yue-PM_B3W3_MACS2_FE.bdg.gz |
| aligned/Bedgraph | loop2-pch2 | 11757-hop1NH-20220125-20220721-20221004-Reps-SK1Yue-PM-PE_B4_W3_MACS2_FE.bdg.gz |
| aligned/Bedgraph | pch2 | 11758-hop1NH-20220125-20220721-20221004-Reps-SK1Yue-PM-PE_B4_W3_MACS2_FE.bdg.gz |
| aligned/Bedgraph | rec8 | Rec8-wildtype-50-75-Reps-SK1Yue-PM_B3W3_MACS2_FE.bdg.gz |
| ReadCounts/txt | wildtype | H2CMNAFX5_n01_Oct_sp7797hop1_S288c_SK1_Yue-PM |
| ReadCounts/txt | loop2 | H2CMNAFX5_n01_Oct_sp11644hop1_S288c_SK1_Yue-PM |
| ReadCounts/txt | loop2-pch2 | H2CMNAFX5_n01_Oct_sp11757hop1_S288c_SK1_Yue-PM |
| ReadCounts/txt | pch2 | H2CMNAFX5_n01_Oct_sp11758hop1_S288c_SK1_Yue-PM |
| ReadCounts/txt | wildtype | HT5W2AFX3_n01_7797e-spike_S288c_SK1_Yue-PM |
| ReadCounts/txt | loop2-pch2 | HT5W2AFX3_n01_11757e-spike_S288c_SK1_Yue-PM |
| ReadCounts/txt | pch2 | HT5W2AFX3_n01_11758e-spike_S288c_SK1_Yue-PM |
| RawSeq/fastq | pch2 | HT5W2AFX3_n02_11758in-spike.fastq.gz |
| RawSeq/fastq | wildtype | HT5W2AFX3_n02_7797e-spike.fastq.gz |
| RawSeq/fastq | wildtype | HT5W2AFX3_n02_7797in-spike.fastq.gz |
| RawSeq/fastq | pch2 | HT5W2AFX3_n02_11758e.fastq.gz |
| RawSeq/fastq | pch2 | HT5W2AFX3_n02_11758e-spike.fastq.gz |
| RawSeq/fastq | pch2 | HT5W2AFX3_n02_11758in.fastq |
| RawSeq/fastq | loop2 | H2CMNAFX5_n01_Oct_11644hop1.fastq.gz |
| RawSeq/fastq | loop2 | H2CMNAFX5_n02_Oct_sp11644in.fastq.gz |
| RawSeq/fastq | loop2 | HKTVNAFX3_n01_11644hop1.fastq.gz |
| RawSeq/fastq | loop2 | H2CMNAFX5_n01_Oct_11644in.fastq.gz |
| RawSeq/fastq | loop2-pch2 | H2CMNAFX5_n02_Oct_sp11757hop1.fastq.gz |
| RawSeq/fastq | loop2 | HKTVNAFX3_n01_11644in.fastq.gz |
| RawSeq/fastq | loop2-pch2 | H2CMNAFX5_n01_Oct_11757hop1.fastq.gz |
| RawSeq/fastq | loop2-pch2 | H2CMNAFX5_n02_Oct_sp11757in.fastq.gz |
| RawSeq/fastq | loop2 | HKTVNAFX3_n02_11644hop1.fastq.gz |
| RawSeq/fastq | loop2-pch2 | H2CMNAFX5_n01_Oct_11757in.fastq.gz |
| RawSeq/fastq | pch2 | H2CMNAFX5_n02_Oct_sp11758hop1.fastq.gz |
| RawSeq/fastq | loop2 | HKTVNAFX3_n02_11644in.fastq.gz |
| RawSeq/fastq | pch2 | H2CMNAFX5_n01_Oct_11758hop1.fastq.gz |
| RawSeq/fastq | pch2 | H2CMNAFX5_n02_Oct_sp11758in.fastq.gz |
| RawSeq/fastq | loop2-pch2 | HT5W2AFX3_n01_11757e.fastq.gz |
| RawSeq/fastq | loop2 | H2CMNAFX5_n01_Oct_sp11644hop1.fastq.gz |
| RawSeq/fastq | wildtype | H2CMNAFX5_n02_Oct_sp7797hop1.fastq.gz |
| RawSeq/fastq | loop2-pch2 | HT5W2AFX3_n01_11757e-spike.fastq.gz |
| RawSeq/fastq | loop2 | H2CMNAFX5_n01_Oct_sp11644in.fastq.gz |
| RawSeq/fastq | wildtype | H2CMNAFX5_n02_Oct_sp7797in.fastq.gz |
| RawSeq/fastq | loop2-pch2 | HT5W2AFX3_n01_11757in.fastq |
| RawSeq/fastq | wildtype | HT5W2AFX3_n01_7797e-spike.fastq.gz |
| RawSeq/fastq | loop2-pch2 | H2CMNAFX5_n01_Oct_sp11757hop1.fastq.gz |
| RawSeq/fastq | loop2-pch2 | HT5W2AFX3_n01_11757in.fastq.gz |
| RawSeq/fastq | wildtype | HT5W2AFX3_n01_7797in-spike.fastq.gz |
| RawSeq/fastq | loop2-pch2 | H2CMNAFX5_n01_Oct_sp11757in.fastq.gz |
| RawSeq/fastq | loop2 | HFL7VAFX3_n01_11644-Hop1NH.fastq.gz |
| RawSeq/fastq | loop2-pch2 | HT5W2AFX3_n01_11757in-spike.fastq.gz |
| RawSeq/fastq | pch2 | H2CMNAFX5_n01_Oct_sp11758hop1.fastq.gz |
| RawSeq/fastq | loop2 | HFL7VAFX3_n01_11644-in.fastq.gz |
| RawSeq/fastq | pch2 | HT5W2AFX3_n01_11758e.fastq |
| RawSeq/fastq | loop2-pch2 | HT5W2AFX3_n02_11757e.fastq |
| RawSeq/fastq | pch2 | H2CMNAFX5_n01_Oct_sp11758in.fastq.gz |
| RawSeq/fastq | loop2-pch2 | HFL7VAFX3_n01_11757-Hop1NH.fastq.gz |
| RawSeq/fastq | pch2 | HT5W2AFX3_n01_11758e.fastq.gz |
| RawSeq/fastq | loop2-pch2 | HT5W2AFX3_n02_11757e.fastq.gz |
| RawSeq/fastq | wildtype | H2CMNAFX5_n01_Oct_sp7797hop1.fastq.gz |
| RawSeq/fastq | loop2-pch2 | HFL7VAFX3_n01_11757-in.fastq.gz |
| RawSeq/fastq | pch2 | HT5W2AFX3_n01_11758e-spike.fastq.gz |
| RawSeq/fastq | loop2-pch2 | HT5W2AFX3_n02_11757e-spike.fastq.gz |
| RawSeq/fastq | wildtype | H2CMNAFX5_n01_Oct_sp7797in.fastq.gz |
| RawSeq/fastq | pch2 | HFL7VAFX3_n01_11758-Hop1NH.fastq.gz |
| RawSeq/fastq | pch2 | HT5W2AFX3_n01_11758in.fastq.gz |
| RawSeq/fastq | loop2-pch2 | HT5W2AFX3_n02_11757in.fastq |
| RawSeq/fastq | pch2 | HFL7VAFX3_n01_11758-in.fastq.gz |
| RawSeq/fastq | pch2 | HT5W2AFX3_n01_11758in-spike.fastq.gz |
| RawSeq/fastq | loop2-pch2 | HT5W2AFX3_n02_11757in.fastq.gz |
| RawSeq/fastq | loop2 | H2CMNAFX5_n02_Oct_11644hop1.fastq.gz |
| RawSeq/fastq | loop2 | HFL7VAFX3_n02_11644-Hop1NH.fastq.gz |
| RawSeq/fastq | loop2-pch2 | HT5W2AFX3_n02_11757in-spike.fastq.gz |
| RawSeq/fastq | loop2 | H2CMNAFX5_n02_Oct_11644in.fastq.gz |
| RawSeq/fastq | loop2 | HFL7VAFX3_n02_11644-in.fastq.gz |
| RawSeq/fastq | pch2 | HT5W2AFX3_n02_11758e.fastq |
| RawSeq/fastq | loop2-pch2 | H2CMNAFX5_n02_Oct_11757hop1.fastq.gz |
| RawSeq/fastq | loop2-pch2 | HFL7VAFX3_n02_11757-Hop1NH.fastq.gz |
| RawSeq/fastq | loop2-pch2 | H2CMNAFX5_n02_Oct_11757in.fastq.gz |
| RawSeq/fastq | loop2-pch2 | HFL7VAFX3_n02_11757-in.fastq.gz |
| RawSeq/fastq | pch2 | H2CMNAFX5_n02_Oct_11758hop1.fastq.gz |
| RawSeq/fastq | pch2 | HFL7VAFX3_n02_11758-Hop1NH.fastq.gz |
| RawSeq/fastq | loop2 | H2CMNAFX5_n02_Oct_sp11644hop1.fastq.gz |
| RawSeq/fastq | pch2 | HFL7VAFX3_n02_11758-in.fastq.gz |
| RawSeq/fastq | pch2 | HT5W2AFX3_n02_11758in.fastq.gz |

**Table S7. Southern Probes**

| Primers | Sequence |
|---|---|
| His4leu2 (probe4) fwd | AGATCTCCTACAATATCATTTTCTCGC |
| His4leu2 (probe 4) rev | ACCGGTGTTGGGCCTTTCAGTG |
| gat1 fowrd | AGCTCAGTGTGCGTTATGCTTCC |
| gat1 rev | GACGAAATACACTAGGCAGG |
| cbp2 fwd | gtg ttc cct cgc tgt aag caa gcg |
| cbp2rev | tgc cca tga agt tct acc tcc gac |

**Table S8. Antibodies used in immunofluorescence analyses**

| Antibody | Host Animal | Working Dilution | Source |
|---|---|---|---|
| Zip1 (yN-16) | Goat | (1/100) | Santa Cruz |
| Hop1 | Rabbit | (1/500) | Nancy Hollingsworth |
| phospho-Hop1 | Rabbit | (1/200) | Andreas Hochwagen |
| Gmc2 | Mouse | (1/500) | Amy MacQueen |
| Donkey anti-goat IgG w/ Cy3 conjugate (1.0 mg) | Donkey | (1/1000) | Jackson |
| Fluorescein (FITC) AffiniPure Donkey Anti Mouse IgG | Donkey | (1/1000) | Jackson |
| Fluorescein (FITC) AffiniPure Donkey Anti Rabbit IgG | Donkey | (1/1000) | Jackson |
| Anti-Rabbit-HRP | Goat | (1/1000) | Kindle Biosciences |
